# Genotyping the self-incompatibility locus of wild and cultivated Brassica using NGS technologies: application to the evaluation of mate limitation in the endangered species *Brassica insularis*

**DOI:** 10.64898/2026.08.02.742276

**Authors:** Sandrine Maurice, Elodie Flaven, Mathieu Genete, Christelle Blassiau, Agnès Mignot, Christophe Petit, Vincent Castric, Xavier Vekemans

## Abstract

1. Self-incompatibility can limit the availability of compatible mates in small and isolated populations, eventually reducing average seed set to the point that the long-term persistence of the populations can be impaired. This phenomenon, named the S-Allee effect, is caused by the loss of alleles (S-alleles) at the self-incompatibility locus (S-locus) due to the intense genetic drift experienced by small populations. Quantifying the diversity of S-alleles is therefore of direct interest for biological conservation, but efficient genotyping methods have been lacking so far because of technical challenges associated with the typically extreme levels of polymorphism and complex genomic structure of the S-locus.
2. We used two alternative approaches to genotype the S-locus using NGS sequencing technologies in four natural populations of the endangered *Brassica insularis* in Corsica. First, we used an NGS amplicon-sequencing approach using generalist primers for each of the two classes of *Brassica* S-alleles. Second, we obtained whole genome shotgun short-read resequencing data and analyzed them with a recently developed bioinformatic pipeline dedicated to hypervariable loci, which we successfully validated on a public dataset comprising 119 cultivated accessions of *B. oleracea*.
3. By combining the two approaches in natural populations of *B. insularis* we identified 31 distinct S-alleles and obtained fully resolved S-locus genotypes for 319 out of 326 sampled individuals. The number of S-alleles varied from four in the smallest population to 18 in the largest one. As a result, the smallest population exhibited very low proportions of compatible individuals, potentially threatening its persistence. We conclude that introducing individuals carrying S-alleles currently absent from the population could help rescue fertility.

## 1. INTRODUCTION

About 45% of flowering plants possesses a self-incompatibility (SI) system, i.e. a genetic system enforcing outcrossing in hermaphrodites (Ferrer et al., 2025). The most common SI systems, homomorphic SI, are usually characterized by the segregation of large series of alleles at the self-incompatibility locus (S-locus). Within the S-locus, each allele (S-allele) is composed of highly linked genes expressed respectively in the pistil and in the pollen or stamen, whose products are interacting to generate self-pollen rejection, or rejection of outcrossing pollen expressing the same specificity (Takayama and Isogai, 2005; Fuji et al., 2016). Within a population, pollen expressing a rare S-allele will be compatible with a larger number of pistils than pollen expressing alleles that are more common, thus enjoying a male fertility reproductive advantage. This negative frequency-dependent selection is a powerful force, and very high numbers of alleles are expected to be maintained at the S-locus within populations (Wright, 1939), a prediction which has indeed been largely confirmed empirically (Lawrence, 2000; Castric and Vekemans, 2004).

Despite this negative frequency-dependent selection acting on the S-locus, small populations subject to strong genetic drift are expected to carry low numbers of S-alleles (Wright, 1939; Schierup et al, 1997), especially if they are highly isolated (Schierup et al., 2000). This reduction in the number of S-alleles will reduce the proportion of compatible plants within populations (*i.e.* mate availability) as plants sharing and expressing the same S-alleles cannot mate successfully (Byers and Meagher, 1992; Vekemans et al., 1998; Busch & Schoen, 2008). Reduction in average mate availability in small populations with a SI system has been indeed documented in many empirical studies using random or diallel cross experiments, e.g. in *Eupatorium perfoliatum* (Byers, 1995), *Brassica insularis* (Glémin et al., 2008), *Ranunculus reptans* (Willi et al., 2005), *Hymenoxys herbacea* (Campbell & Husband, 2007), *Rutidosis leptorrhynchoides* (Young & Pickup, 2010). Some exceptions occur, however, and two different reasons have been invoked to explain a lack of relationship between population size and mate availability in some studies. First, the type of SI system, as theoretical investigations have shown that limitations in mate availability are expected to be stronger in sporophytic SI systems, i.e. systems in which the pollen phenotype is encoded by the diploid paternal genotype, than in gametophytic systems, i.e. systems in which the pollen phenotype is determined by its own haploid S-locus genotype (Vekemans et al., 1998; Levin et al., 2009). This is mainly because pollen from species with gametophytic SI segregate for two different S-alleles so that half of the pollen of plants sharing a single S-allele are cross-compatible (Hoebee et al. 2012). The second reason invoked is associated with differences in genetic connectivity among populations. Indeed, even moderate levels of migration can impede the loss of S-allele diversity within small populations because negative frequency-dependent selection is causing high effective gene flow at the S-locus (Schierup et al. 2000). This has been suggested to cause the patterns observed in *Leavenworthia alabamica*, a species with a SSI system, where relatively high allelic diversity (> 20 S-alleles) and high mate availability (> 80%) were reported independently of large variation in population size (Busch et al., 2010).

If the number of S-alleles becomes too small, many individuals will share the same S-alleles and even pollinated plants can fail to produce seeds by lack of successful fertilization, resulting in a mate-finding Allee effect (Gascoigne et al., 2009) that is specific of species with functional SI, a phenomenon that has been called the S-Allee effect (Wagenius et al., 2007; Leducq et al., 2010). Direct proof of the S-Allee effect was reported in two species with SSI, where the diversity of S-alleles was experimentally manipulated (Elam et al. 2007, Leducq et al., 2010), and a key prediction of the S-Allee effect (a reduction of seed-set in small populations) was verified in several self-incompatible species (e.g. Widén, 1993; Reinartz & Les, 1994; Morgan, 1999; Honnay et al., 2006), although these studies lacked direct estimations of S-allele diversity.

Current census size is an imperfect predictor of the diversity of S-alleles in a population, which also depends on population history and genetic isolation (Schierup et al., 2000), so a proper evaluation of the severity of the S-Allee effect in nature requires direct quantification of allelic diversity at the S-locus. This has typically been achieved either through diallel crosses or through series of random tests of pairwise compatibility, but conducted on very small samples, far from the population-scale resolution that is required.

Genes controlling SI have been identified in only nine plant families so far (Zhang et al. 2024), and despite the genomic revolution driven by next-generation sequencing technologies, molecular characterization of S-allele diversity remains scarce in non-model species. This limited progress reflects the intrinsic difficulty of genotyping the S-locus in species with multiallelic SI (Jorgensen et al. 2012), since strong balancing selection maintains extensive allelic diversity and exceptionally high sequence divergence among S-alleles (Vekemans & Slatkin, 1994). As a result, PCR-based methods have consistently faced the difficulty of designing primers that amplify all S-alleles of a given species, and shotgun or RNAseq short-read resequencing methods have been limited by the difficulty of mapping short-reads to single reference sequences (Jorgensen et al. 2012; Vekemans et al., 2021). In addition, paralogues of the S-locus genes have been reported, with genomic locations either outside (e.g. in *Arabidopsis lyrata*, Charlesworth et al., 2003a) or inside the S-locus (*e.g.* in Brassica species, Kusaba et al., 1997). Some of these paralogues have strong phylogenetic proximity with actual S-alleles, further complicating S-locus molecular genotyping (Charlesworth et al., 2003b; Kusaba et al., 1997).

Several approaches have been proposed in the last decade to resolve the issues associated with the use of next generation sequencing technologies in genotyping highly polymorphic loci such as the S-locus (Vekemans et al., 2021). These approaches target either transcriptomic data, obtained with short-read RNAseq (e.g. Ramanauskas et al., 2025) or long-read RNAseq methods (e.g. Badouin et al., 2021; Maenosono et al., 2024), or genomic data. For the latter, the approaches proposed are based either on PCR with general primers followed by short-read amplicon sequencing (e.g. Jorgensen et al. 2012) and data analyses using clustering techniques, or on whole genome shotgun short-read sequencing followed by data analyses using mapping on multiple references (De Franceschi et al., 2018; Genete et al., 2020), either on the full dataset or after prior filtering of reads using a k-mer library (Genete et al., 2020), or directly by searching for allele-specific k-mers (Cai et al., 2026).

*Brassica insularis* is an endangered species endemic to Corsica and Sardinia, with most populations showing a severe decline (Noël et al., 2010). This self-incompatible species expresses a SSI system homologous to that of *B. oleracea* and *B. rapa,* which has been extensively characterized at the molecular and genomic levels (e.g., Takayama & Isogai, 2005; Cui et al., 2020; Goring et al., 2023). A previous study estimated that the number of S-alleles ranges from only five in a small population (N ≈ 100) to 19 in a larger population (N ≈ 1000–2000; Glémin et al., 2005, lowest estimates). *In situ* pollination success and fruit-set data further suggested that reproduction may be negatively affected by low S-allele diversity in the smallest population (Glémin et al., 2008). However, S-allele diversity was estimated based on a crude biochemical assay of stylar protein recognition, which did not allow precise identification of S-alleles. Several genotypes could not be fully resolved, and missing S-alleles accounted for up to 50% of all sampled gene copies, generating substantial uncertainty in both S-allele diversity estimates and individual genotype assignments, from which mate-availability estimates were inferred (Glémin et al., 2008). In the cultivated *Brassica oleracea* and *B. rapa*, S-allele diversity has historically been characterized using PCR amplification of DNA or cDNA with general primers, followed by cloning and sequencing (e.g. Watanabe et al., 2000; Sato et al., 2002). To date, sequences are available for nearly 50 alleles in each species (Yamamoto et al., 2023). Several strategies have been developed to genotype individuals at the S-locus for breeding purposes, including PCR-RFLP (Nishio et al., 1996; Park et al., 2002), dot-blot analyses with allele-specific oligonucleotide probes targeting the highly polymorphic pollen gene *SCR/SP11* (Oikawa et al., 2011) or hypervariable regions of the pistil gene *SRK* (Haseyama et al., 2018), and PCR amplification with allele-specific primers (Oikawa et al., 2011).

Here, we leverage recent advances in sequencing technologies to obtain a more accurate estimate of S-allele number in *B. insularis* and to maximize the recovery of complete individual genotypes at the S-locus. We tested two alternative approaches using NGS sequencing technologies to genotype the S-locus on large samples from the same four natural populations of *Brassica insularis* as those used by Glémin et al. (2008). First, we used a PCR-based approach using a combination of generalist primers, followed by short-read amplicon sequencing. Second, we performed whole genome shotgun resequencing using short-read sequencing, followed by application of a bioinformatic pipeline dedicated to hypervariable loci (Genete et al., 2020) using previously known S-allele sequences from cultivated Brassica as references. To validate the pipeline, we analyzed a public dataset comprising 119 cultivated accessions of *B. oleracea*. We then compared the relative performances of the two approaches, quantified the diversity of S-alleles found in each population and compared it with previous estimates and with theoretical predictions, and finally used these S-locus genotypes to calculate expected mean and individual-based mate availability to determine the extent to which these populations may be suffering from an S-Allee effect.

## 2 MATERIALS AND METHODS

### 2.1 Population sampling

*Brassica insularis* is a rare Mediterranean species known from a few populations in Corsica, Sardinia and North Africa (Snogerup et al., 1990). It is a long-lived species that grows on limestone or schist cliffs. It is pollinated by insects and presents a sporophytic SI system. It is a close relative of the cultivated cabbage *Brassica oleracea*, with 2*n*=18 chromosomes (Saban et al., 2023).

We sampled four populations of various sizes across Corsica that are monitored for demography (Noël et al., 2010; Fig. 1). Approximate population sizes at the beginning of the 21^th^ century were estimated to be 2,000 for Teghime (TG), 500 for Inzecca (IZ), 300 for Punta Corbaghiola (CG), 80 for Punta Calcina (CA) by Glémin et al. (2005). TG, the largest population, is stable, while the three others are declining and have a high extinction probability at 50 years (Noël et al., 2010). In CA, the whole population is monitored; in CG and IZ the intermediate-size populations, an area containing all accessible plants is monitored; and in TG, only a subset of the population is monitored, which consists in two cliffs separated by 260 meters. In the areas studied, we collected in 2014 a leaf sample of all accessible plants that had flowered between 2011 and 2014. Subsequently, new flowering plants were also sampled until 2022. Although sampled plants were generally well separated, in a few cases—particularly in TG, where local density is high—some physically distinct shoots (ramets) collected may belong to the same genet. This could not be verified in the field because the root systems extend deeply into the rock substrate. For this study, we obtained leaf tissue from 326 individuals: 44 individuals from CA; 48 from CG; 74 from IZ; and 160 from TG.

**Figure 1.**
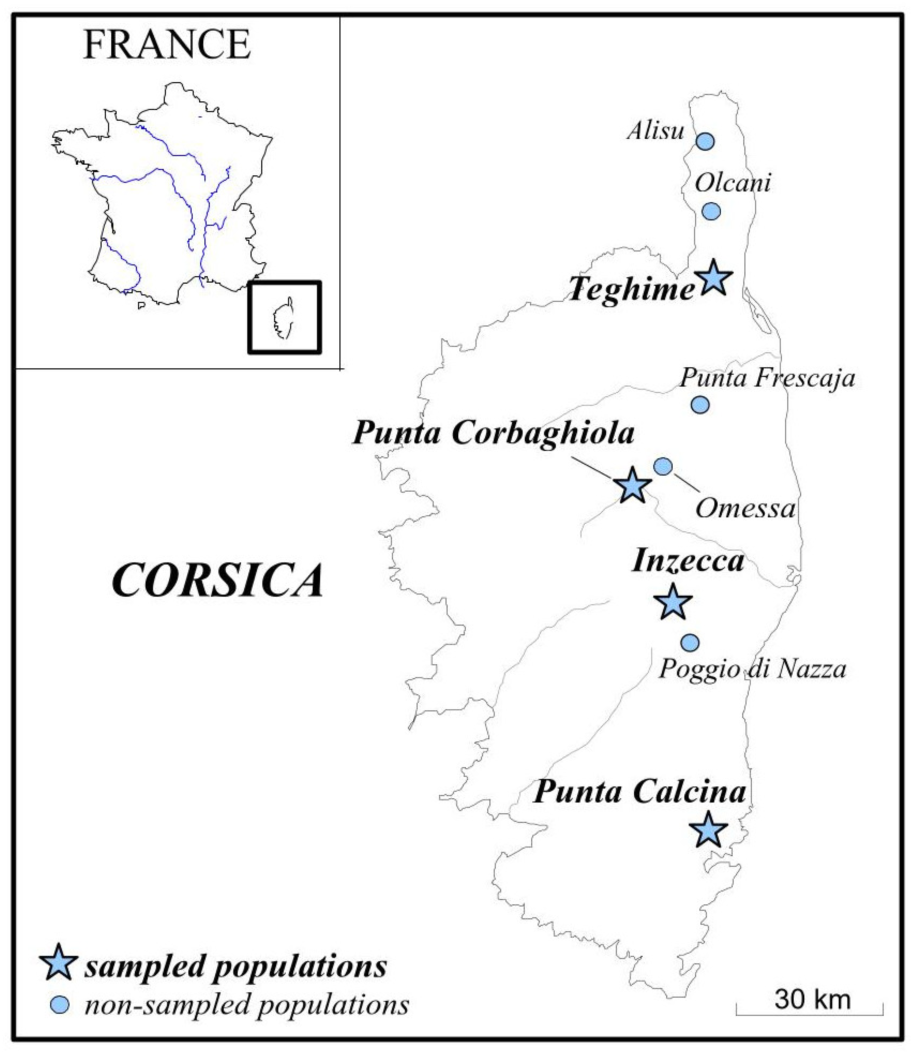
Localization of the four studied populations: Teghime (TG), Punta Corbaghiola (CG), Inzecca (IZ), and Punta Calcina (CA).

### 2.2 Genotyping the S-locus with the amplicon sequencing approach

In 2014, total DNA was extracted from leaf tissue using the CTAB method (Doyle and Doyle, 1987). From 2015 onwards, we used the Perkin Elmer chemagic DNA Plant Kit and the Kingfisher flex automated extraction system (Thermo Fisher Scientific). Based on allelic sequences of the S-locus pistil gene *SRK* (*S-locus Receptor Kinase*) of *B. oleracea* obtained from Genbank (Yamamoto et al., 2023), we designed general forward primers separately for the two phylogenetic classes of alleles (class I and class II) that target an amplicon size around 380 nucleotides (class1F: ^5’^ TCACCAGTGCGATATGTACA^3’^, class2F: ^5’^ ACGTGTGCGATCCGCTTTAC ^3’^) when combined with the universal reverse primer SLGR (Mable et al., 2003; SLGR: ^5’^ ATCTGACATAAAGATCTTGACC ^3’^).

We first sequenced amplicons with the Sanger method. For this, PCRs were carried out in 20 µl final volume including 0.5 µM of each primer and 1 µl of genomic DNA, using the Master MIX Phusion 2X kit (Thermo Fischer Scientific). The cycling scheme was 2 min at 98°C, 30 cycles including 30 s initial denaturation at 98°C, 60 s annealing at 58°C (class II) or at 56°C (class I), and 30 s extension at 72°C, and then 15 min final extension at 72°C. Sanger sequencing was performed on the PCR products by the Eurofins MWG operon sequencing platform (F-Cochin) and sequences were analyzed with the Geneious 11.1.5 software. 162 individuals were analyzed using this method. As some genotypes could not be solved despite positive amplifications, possibly because of cross amplification between the *SRK* and *SLG* paralogs or because of the presence of two alleles of the same class, we turned to short-read next-generation sequencing.

For amplicon NGS sequencing, PCRs were carried out following the same protocol as before except that we included only 0.25 µM of each primer. Sequencing was performed on a MiSeq sequencer using kits developed by Illumina (Illumina 2X250 bp pair-end sequencing kit) at the CEMEB-LABEX platform. In total 238 and 230 samples were sent to be sequenced after amplification with class II and class I primers, respectively. To discriminate among paralogous and allelic *SRK* sequences, we analyzed sequencing data with the AMPLISAS pipeline (Sebastian & al, 2016), which includes a phase of clustering of similar sequences before filtering them by frequency. Final parameters used were default settings except for the frequency threshold under which sequences are discarded (“minimum per amplicon frequency”) that was set to 4%. The pipeline outputs the number of reads for each cluster and its consensus sequence. We then used the sequence similarity search tool YASS (Noe & Kucherov, 2005) to compare these sequences with reference *SRK* allele sequences and with *SRK* paralogs from *Brassica oleracea* and *B. rapa* (available at https://doi.org/10.48579/PRO/EUP5ZX), including *SLG* which co-segregates with *SRK* at the S-locus (Nasrallah et al., 1991).

### 2.3 Genotyping the S-locus with the whole-genome resequencing approach

The amplicon sequencing (AS) approach could not resolve all S-locus genotypes, partly because of null S-alleles (*SRK* sequences that cannot be amplified with the set of primers), and partly because of allelic competition in the PCR amplifications. Hence, we used in parallel a genotyping method based on bioinformatic analyses of whole-genome resequencing data (Genete et al. 2020). To test the method, we first analyzed published whole-genome resequencing data from 119 accessions of cultivated *Brassica oleracea* (Cheng et al., 2016). The dataset was analyzed with NGSgenotyp (Genete et al. 2020), a bioinformatic pipeline that maps sequentially individual short reads against reference S-locus sequences and computes mapping statistics in order to determine the corresponding S-locus genotype of each individual. To run NGSgenotyp, we built a reference library of *SRK* and *SLG* sequences (available at https://doi.org/10.48579/PRO/EUP5ZX) from *B. oleracea* (41 *SRK* and 40 *SLG*) and *B. rapa* (36 *SRK* and 41 *SLG*) from Genbank, based on information from Yamamoto et al. (2023). Because of strong cross-mapping of reads on both *SRK* and *SLG* sequences of the same S-haplotype, we modified the pipeline by computing additional mapping statistics counting only the reads with zero mismatch with the reference sequence (https://gitlab.univ-lille.fr/eep_lab/mathieu_genete/NGSgenotyp). As some ambiguities associated with the confusion between *SRK* and *SLG* remained using this approach, we also applied the NGSgenotyp pipeline targeting the *SCR* (pollen) gene (with the k-mer size for filtering set to 15 instead of the default value of 20, and by reporting mapping statistics computed on local alignment, see Duan et al. 2024; command line: NGSgenotyp genotyp -ks 15 -k -A local -M 0 -o out_folder_path -i read_list_filename -d RefSeq_filename.fa), using a reference library of *SCR* exon 2 sequences from *B. oleracea* (41 sequences) and *B. rapa* (45 sequences) obtained from Genbank (available at https://doi.org/10.48579/PRO/EUP5ZX). We focused on exon 2 of *SCR* because it contains the sequence of the excreted SCR peptide. This approach using *SCR* sequences was very powerful and allowed to resolve ambiguous genotypes.

In *B. insularis*, we obtained original short-read whole genome resequencing data for 246 individuals. We performed three sequencing experiments, respectively in 2021 (19 individuals), 2022 (198 individuals) and 2024 (38 individuals, including 9 duplicates from the 2021 or 2022 experiments). For the 2021 experiment, we purified DNA from 15 mg of dried leaves of each sample with chemagic beads (PerkinElmer), following Holtz et al. (2016), using the manufacturer’s instructions but with an additional Agencourt AMPure beads (Beckman) purification. In the 2022 and 2024 experiments, we purified DNA from 15 mg of dried leaves of each sample with nucleomag 96 plant (Macherey Nagel). DNA was then quantified by Qubit with the DNA HS kit. In 2021, 50 ng of DNA was fragmented mechanically with Bioruptor (Diagenode) to obtain fragments of around 200-300 bp, which we verified using a BioAnalyzer (Agilent) with a DNA HS chip. In 2022 and 2024, 50 ng of DNA was fragmented enzymatically with the Nextflex kit. We then prepared indexed genomic libraries with the Nextflex Rapid DNA Seq kit V2.0 (PerkinElmer) in 2021, with the Nextflex Rapid xp DNA seq kit 2.0 in 2022, and with the Nextflex Rapid xp DNA seq kit 2.0 V2 in 20224, using manufacturer’s instructions. Briefly, extremities of sequences were repaired and tailed, ligated with universal adaptors P5/P7 containing a multiplexing unique dual index (PerkinElmer), followed by amplification with five or six cycles of PCR. We then selected fragments between 150 and 300 bp with AMPures beads and pooled all libraries within each experiment in equimolar proportions. The prepared libraries were sequenced by Illumina NovaSeq (2x 150pb, paired-end) at the GenoScreen platform (Lille, France, in 2021 and 2024) or at the Ligan platform (Lille, France, in 2022).

The raw sequencing data was analyzed with NGSgenotyp separately on *SCR* and *SRK*, using the same reference libraries of *B. oleracea* and *B. rapa* sequences as described above. After one round of analysis, the libraries were enriched with new *B. insularis SCR* and *SRK* sequences obtained using the *de novo* assembly module of NGSgenotyp, and the genotyping procedure was resumed.

### 2.4 Assembling datasets produced by the two approaches and allele naming scheme

Overall, about 56% (*N*=182) of individuals were analyzed with both methods, including all individuals with an unresolved genotype with the AS approach for which enough plant material was available. A final dataset was then assembled by combining all individuals genotyped with the WGS approach, with those analyzed only by the AS approach.

To name the S-alleles identified in *B. insularis* we adopted the following approach. We determined trans-specifically shared alleles between *B. insularis* and *B. oleracea* based on the proportion of pairwise nucleotide differences (*p*) at exon 2 of the *SCR* gene, with a cut-off value at *p* = 0.025. Indeed, the distribution of this statistic was found to be highly bimodal, with 21 interspecific pairs showing values varying between 0 and 0.021 (mean *p* = 0.006), whereas all other pairs showed values varying between 0.13 and 0.62 (mean *p* = 0.44; Fig. S3). For these S-alleles shared trans-specifically with *B. oleracea*, we applied the same ID *in B. insularis* as in *B. oleracea* (e.g. BiS06 for the allele trans-specific to allele S06 from *B. oleracea*, Fig. 2). For S-alleles that are not shared with *B. oleracea*, we labelled them arbitrarily, respectively BiS85, BiS90, BiS92, BiS93, BiS94, BiS97 and BiS102. In this manuscript, we also introduce a naming system for functional S-alleles in *Brassica* inspired from that used for *Arabidopsis* and *Capsella* (Duan et al. 2024). We use a 4-digit system with the first digit corresponding to the allelic class (1 for class I alleles, 2 for class II alleles), and the last three corresponding to a label shared by all trans-specific alleles with the same functional specificity. As the largest number of S-alleles described today in the genus Brassica correspond to *B. oleracea*, we used the number of the S-allele in *B. oleracea* as reference ID, e.g. the functional specificity of allele S06 from *B. oleracea* will be named H1006 (note that the H is for functional Haplotype) because it is a class I allele.

**Figure 2.**
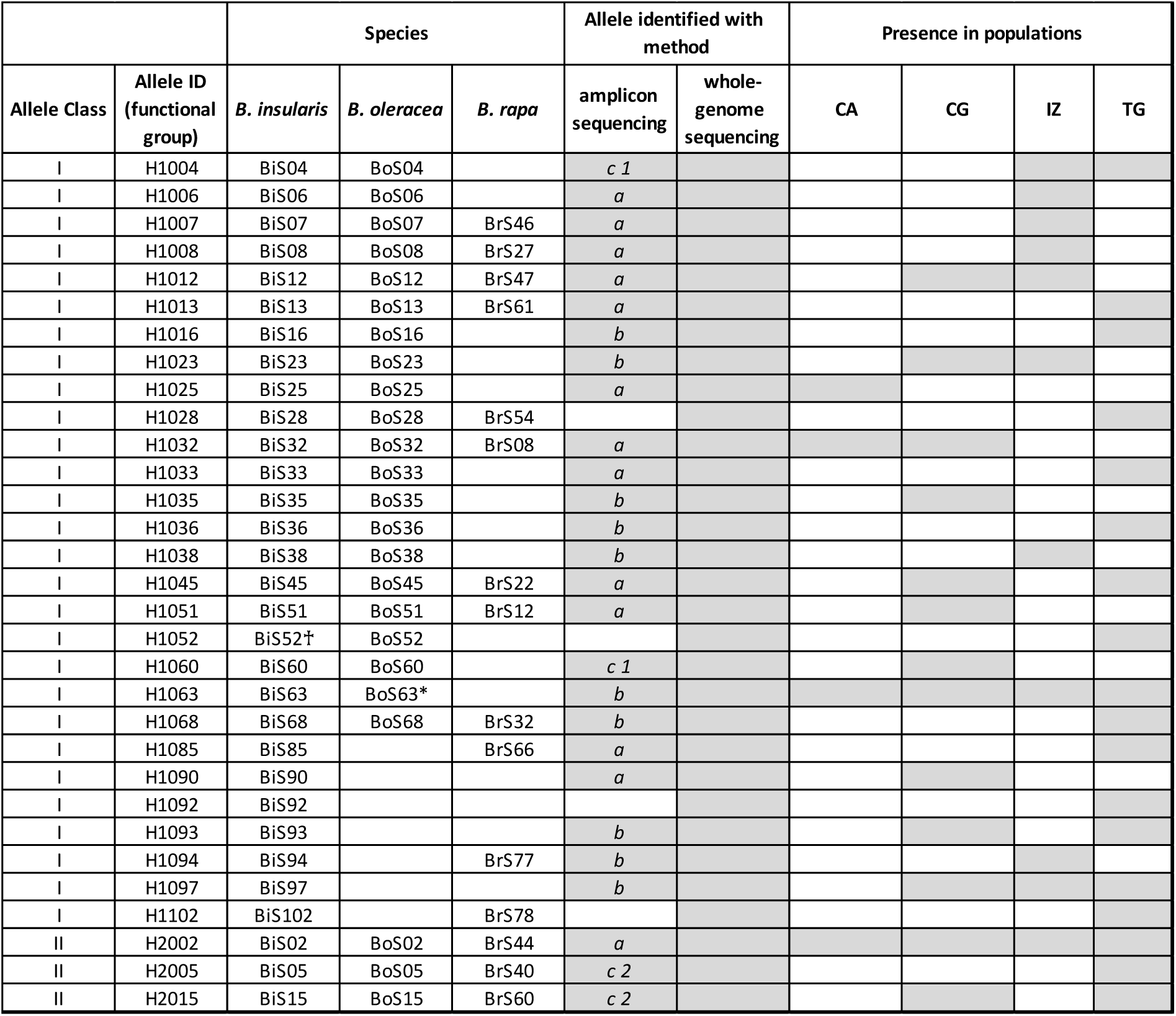
List of S-alleles identified through AS and WGS approaches in *Brassica insularis*. Presence of each S-allele in each population is indicated. For the AS column: “a” means the allele was always revealed; “b” means the allele was only sometimes revealed; “c” means the allele was not distinguished from another allele (bearing the same number 1 or 2). The *B. oleracea* allele marked with * is missing a *SCR* sequence so that its relationship with the corresponding *B. insularis* allele was determined based on similarities of *SRK* sequences. The *B. insularis* allele marked with ♰ is putatively non-functional as the *SCR* sequence carries a one-bp deletion in exon 2.

### 2.5 Allelic diversity per population and mate availability estimation

Allelic frequencies and population differentiation (F_ST_ statistic) were calculated using the R package genepop (Rousset, 2008). To compute average mate availability within population, i.e. the average proportion of individuals in the population compatible to a randomly chosen individual (Vekemans et al. 1998), we used the dominance relationships between alleles known from *Brassica* (Thompson & Taylor, 1966; Hatakeyama et al. 1998; Kakizaki et al., 2003) to define compatibility between individuals, with some simplification. In the pistil, all alleles were assumed to be codominant. In pollen, all alleles of class I were assumed to be dominant over alleles of class II; codominance was assumed among all alleles of class I and dominance among class II alleles were defined according to the H2002>H2015>H2005 relationship (i.e., BoS02=BrS44>BoS15=BrS60>BoS05=BrS40; Kakizaki et al., 2003). Because of asymmetries between dominance in pollen and pistil, the number of compatible partners of an individual may differ between its male and female functions (Vekemans et al., 1998). We here report the proportion of compatible partners from the female function perspective, computed as the proportion of individuals in the population that can act as compatible pollen donors for a given maternal individual, as we are interested in a possible limitation of seed production in the wild.

### 2.6 Numerical simulations to estimate expected allelic diversity in small populations

We used the simulation program of sporophytic self-incompatibility from Schierup et al. (1997) to obtain estimates of expected S-allele diversity in populations with sizes corresponding to the *B. insularis* populations investigated here. We investigated two of the models proposed by Schierup et al. (1997), representing contrasted situations regarding patterns of dominance among S-alleles: 1) a model with strict codominance in the determination of the pollen and pistil SI phenotypes (SSI-COD) and 2) a model with strictly hierarchical dominance relationships among alleles in pollen and codominance in pistil (SSI-DOMCOD). As described in the previous section, the observed patterns of dominance relationship in Brassica are intermediate between these two models. As evidence for pollen limitation of reproductive output has been reported in *B. insularis* (Glémin et al., 2008), we implemented strong pollen limitation in the simulations by discarding a potential maternal parent if the first pollen source sampled from the population was found to be incompatible (Vekemans et al., 1998). We simulated populations with sizes corresponding to empirical estimates from *B. insularis*, i.e. *N* = 80, 300, 500, and 2000, and used a mutation rate to new S-alleles of 1. 10^-6^. Each simulation was started with 2*N* functionally different alleles in the population and allowed to evolve for 100,000 generations, a time chosen to ensure that mutation–selection–drift equilibrium was attained. We recorded the number of S-alleles present, and replicated the whole process 100 times for each value of *N* and each dominance model. For comparison with species with gametophytic SI, we also computed for each case the expected allelic diversity using the analytical formula derived by Yokoyama & Nei (1979).

## 3 RESULTS

### 3.1 Evaluating the amplicon sequencing approach to genotype the S-locus

We amplified DNA separately with two pairs of generalist PCR primers targeting class I and class II alleles of the S-locus pistil gene *SRK* from Brassica. Sanger sequencing revealed the unexpected presence of class II *SRK*-like sequences in the majority of individuals, which would represent an abnormally high frequency of class II alleles. Inspection of these sequences revealed that they correspond to an unlinked paralog of *SRK* (*SLR2*; Nasrallah et al., 1991) that was co-amplified with the class II primers. Hence, we replaced the Sanger sequencing by an amplicon NGS sequencing approach using Illumina MiSeq technology, followed by sequence separation using the AMPLISAS clustering pipeline (Sebastian et al, 2016). Overall, 262 individuals were analyzed with this approach.

Amplicons obtained with the class II primers led to the identification of three groups of *SRK*-like sequences: one group with high similarity to the *SLR2* paralog, that we discarded; one similar to allele BoS02 from *B. oleracea* and one similar to allele BoS05. We thus identified two class II S-alleles, that we named, respectively, BiS02 and BiS05. With PCR primers designed to amplify class I sequences, we detected 28 *SRK*-like sequences. Ten of them always appeared as pairs, as is typical of the common linkage of *SRK* with its closest paralogue, *SLG,* in Brassica species (Nasrallah et al., 1991), and we interpret these five pairs as indicating the segregation of five individual S-alleles. For 18 of the sequences, we could identify a highly similar sequence (i.e. less than 5% nucleotide divergence; Fig. 2) from *B. oleracea* (labelled either as *SLG* or *SRK* in that species). In addition, we found five sequences with higher divergence to any known *B. oleracea* S-allele (nucleotide divergence from the closest *B. oleracea* allele ranging from 9 to 15%), suggesting that they represent five S-alleles that were not currently identified in *B. oleracea*.

Combining results from class I and class II sequencing, we identified 25 S-alleles in *B. insularis*. We could not identify any S-allele for five individuals. We identified 101 heterozygous individuals (with 82 inter-classes heterozygotes and 19 intra-classes heterozygotes; Table 1.a). For 156 individuals, we identified only one S-allele (125 had only a class I allele and 31 only a class II allele). Homozygotes for a class I allele should not be produced with a functional SI system, as class I alleles are co-dominant with one another and dominant over class II alleles, so individuals where we detected only one class I allele likely indicate incompletely resolved genotypes. In contrast, individuals where we detected only one class II allele could be true class II homozygotes, or incompletely resolved genotypes. Overall, this genotyping method based on PCR amplicons led to a proportion of missing gene copies between 25.8 and 31.7%.

**Table 1:** Numbers of individuals analyzed by each method and resulting genotypic classes. In table b. numbers in brackets represent the number of individuals analyzed by both methods, in table c. numbers in brackets give the number of individuals with fully resolved genotypes. Missing genes are represented by a point. In AS method, a CII/. genotype could be a class II homozygote or a CII/. heterozygote. In WGS method, CII/. corresponds to a CII homozygote (see method). For both method, CII/CII corresponds to CII heterozygotes. All CI/CI are heterozygotes.

a. Amplicon Sequencing
| Population | Nb ind | ./. | CI/. | CII/. | CI/CI | CI/CII | CII/CII |
| --- | --- | --- | --- | --- | --- | --- | --- |
| CA | 41 | 0 | 5 | 11 | 0 | 25 | 0 |
| CG | 30 | 0 | 6 | 0 | 8 | 15 | 1 |
| IZ | 58 | 1 | 26 | 6 | 3 | 22 | 0 |
| TG | 133 | 4 | 88 | 14 | 7 | 20 | 0 |
| Total | 262 | 5 | 125 | 31 | 18 | 82 | 1 |

b. Whole Genome Sequencing
| Population | Nb ind<br>(common) | ./. | CI/. | CII homo | CI/CI | CI/CII | CII/CII |
| --- | --- | --- | --- | --- | --- | --- | --- |
| CA | 22 (19) | 0 | 0 | 9 | 9 | 4 | 0 |
| CG | 33 (15) | 0 | 1 | 0 | 12 | 17 | 3 |
| IZ | 55 (39) | 0 | 0 | 4 | 40 | 11 | 0 |
| TG | 136 (109) | 1 | 0 | 0 | 88 | 41 | 6 |
| Total | 246 (182) | 1 | 1 | 13 | 149 | 73 | 9 |

c. Both methods combined
| Population | Nb ind |  |  |  |  |  |  |
| --- | --- | --- | --- | --- | --- | --- | --- |
|  | (complete) | ./. | CI/. | CII homo | CI/CI | CI/CII | CII/CII |
| CA | 44 (44) | 0 | 0 | 10 | 5 | 29 | 0 |
| CG | 48 (47) | 0 | 1 | 0 | 17 | 27 | 3 |
| IZ | 74 (74) | 0 | 0 | 4 | 43 | 27 | 0 |
| TG | 160 (154) | 1 | 5 | 0 | 94 | 54 | 6 |
| Total | 326 (319) | 1 | 6 | 14 | 159 | 136 | 9 |

### 3.2 Genotyping the Brassica S-locus using whole genome resequencing data

We then evaluated whole-genome resequencing approaches using the NGSgenotyp pipeline (Genete et al., 2020) to genotype the S-locus in 119 accessions of *Brassica oleracea*. To obtain reliable genotyping results, with very limited cross-mapping among alleles or paralogs and clear discrimination between homozygous and heterozygous genotypes, we target the S-locus gene expressed in anthers (*SCR,* also called *SP11*) rather than *SRK*, using a reference database of *SCR* exon 2 sequences from *B. oleracea* and *B. rapa*. Using the *de novo* assembly module of NGSgenotyp, we identified 29 distinct putative *SCR* sequences, including three class II and 26 class I alleles. Among these, 24 closely matched sequences already present in the *B. oleracea* database, and five corresponded to newly discovered *SCR* sequences. By subsequently applying the NGSgenotyp pipeline to *SRK* in individuals carrying these novel *SCR* sequences, we associated them with five known *B. oleracea* S-alleles lacking *SCR* sequences in GenBank (BoS23, BoS36, BoS38, BoS50, and BoS60). After updating the reference database, we reran the NGSgenotyp pipeline on *SCR*, using *SRK* genotyping for confirmation. This yielded complete S-locus genotypes for 116 of the 119 accessions (Table S1). Two additional accessions (Broccoli_11 and Broccoli_12) showed three S-alleles, potentially reflecting sample contamination or polyploidy. In the remaining accession (Kohlrabi_6), neither *SCR* nor *SRK* reads were detected despite normal coverage at control genes (∼10×), possibly suggesting deletion of the S-locus. Among the 116 successfully genotyped accessions, five lacked detectable *SCR* signal, whereas *SRK* of these accessions unambiguously identified homozygous genotypes for either allele BoS16 (Kohlrabi_9, Kohlrabi_13, Kohlrabi_14, and Cauliflower_17) or BoS28 (Chinese_kale_2), so that the total number of S-alleles detected is 31 (Table S2). For BoS28, other accessions carrying the same allele displayed normal *SCR* signal, suggesting deletion of the *SCR* gene in accession Chinese_kale_2. The four BoS16 accessions were the only representatives of this S-haplotype in the dataset, and both *SCR* and *SRK* were absent, whereas *SLG* was still detected, again possibly suggesting structural alteration of these S-haplotypes. Overall, 104 accessions out of 116 were homozygous and 12 (10%) heterozygous at the S-locus. Of the 232 S-allele copies identified, 52% belonged to class II and 48% to class I. The two most frequent alleles were BoS15 and BoS02, both belonging to class II (Fig. S1).

We applied the NGSgenotyp pipeline on the whole genome resequencing short Illumina reads generated for 246 *B. insularis* individuals (average depth: 13X) using the database of reference *SCR* sequences obtained for *B. oleracea*, including the newly discovered *SCR* sequences described above. This resulted in the identification of 31 putative S-alleles in *B. insularis* (28 class I alleles and 3 class II alleles; Fig. 2 & Table S3). Twenty-three of them were trans-specifically shared with *B. oleracea* (with a proportion of pairwise nucleotide differences lower than 0.025, see Material & Methods). Three S-alleles were trans-specifically shared with *B. rapa* (according to the same criterion) but absent from *B. oleracea.* The five other *SCR* sequences we identified were more distant from any other known *SCR* of *B. oleracea* or *B. rapa* (with a proportion of nucleotide differences to the closest interspecific allele ranging from 0.15 to 0.33) and may represent newly discovered S-alleles in Brassica. We analyzed partial *SRK* sequences obtained from the individuals carrying these five newly discovered *SCR* sequences and found a closely related *B. oleracea SRK* sequence (with a proportion of nucleotide differences <0.007) for one of them (BoS63; Fig. 2). For individuals carrying the four remaining unknown *SCR* sequences, we could not identify *SRK* sequences that are closely related to sequences from the other Brassica species, suggesting that they correspond to entirely new Brassica S-alleles (respectively named BiS90, BiS92, BiS93, and BiS97).

Full or partial *SCR* exon 2 and *SRK* exon 1 sequences were obtained for all 31 S-alleles (available at https://doi.org/10.48579/PRO/EUP5ZX). For shared alleles, nucleotide identities for *SCR* and *SRK* were always higher when compared with *B. oleracea* than with *B. rapa* alleles (Fig.3, Fig.S2), in agreement with the species phylogenetic relationships (Saban et al., 2023). By combining *SCR* and *SRK* genotyping procedures on the whole genome resequencing data, we obtained fully resolved genotypes for 244 of the 246 individuals sequenced (Table 1.b). The two remaining individuals showed insufficient sequencing depth for complete genotyping.

**Figure 3.**
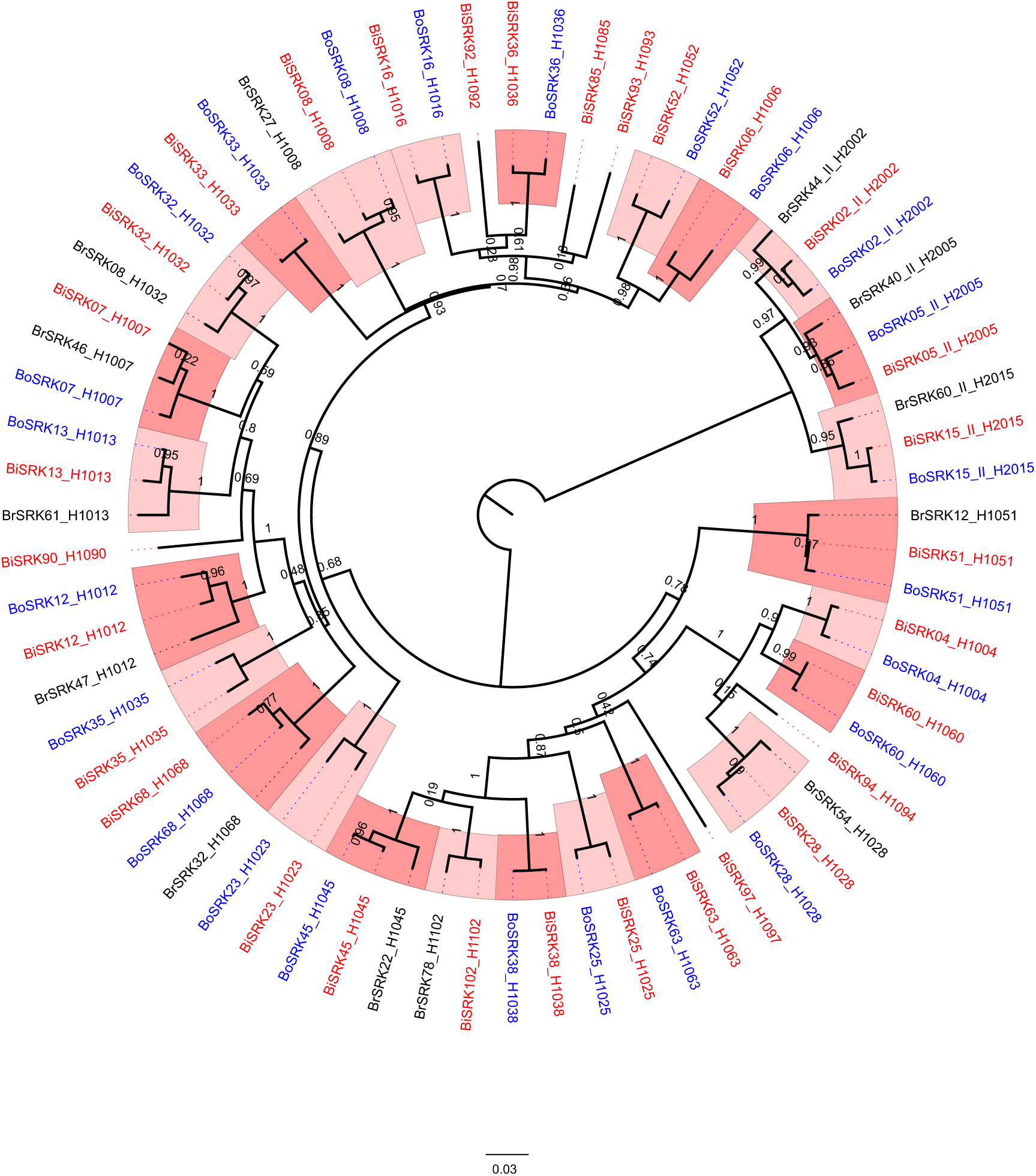
Phylogenies of *SRK* sequences from *Brassica insularis* (Bi, in red), *B. oleracea* (Bo, in blue) and *B. rapa* (Br, in black), as obtained using tree reconstruction based on maximum likelihood with PhyML. Node support obtained from bootstraps is indicated. Groups of trans-specifically shared S-alleles are represented by alternate pink shades and labeled as HXYYY where X indicates whether the allele belongs to class I (1) or II (2), and YYY represents the label of the functional group.

### 3.3 Combining results from the two methods

While 262 individuals were analyzed with amplicon sequencing (AS) and 246 with the whole genome sequencing (WGS) methods, 182 individuals were analyzed by both methods. This allowed us to evaluate their congruence based on similarity of S-allele sequences and on the composition of individual genotypes. Whenever one or two alleles were revealed by AS in an individual, genotyping with WGS always confirmed the presence of these alleles, with a single exception for which we suspect a possible label switching or DNA contamination. Conversely, alleles revealed by WGS and not by AS fell in two categories (Fig. 2): alleles that were never revealed by AS (four alleles in total: BiS28, BiS52, BiS92, BiS102), and alleles that were only partially retrieved with AS (i.e. only in some individuals). This allowed us to resolve all incomplete or ambiguous genotypes obtained by the AS. Moreover, we found two sets of sequences that had been grouped together in the AS method but were in fact composed of two closely related but different S-alleles each, as revealed by results from the WGS method (sets c1 and c2 in Fig.2). First, among the three sequences obtained with AS close to allele BoS05 from *B. oleracea*, two, that always came together, were confirmed by WGS to correspond to *SRK* and *SLG* of *B. oleracea* BoS05, while the third one was shown to actually correspond to *SRK* of BoS15. Second, a set of two sequences found with AS to fall within the cluster of BoS04 and BoS60 from *B. oleracea* was confirmed to correspond to either BoS04 or BoS60, depending on the individuals.

Overall, by combining the two datasets we were able to obtain fully resolved genotypes for 319 out of 326 individuals tested (Table 1.c & Table S4), including 44 individuals from CA, 47 from CG, 74 from IZ, and 160 from TG.

### 3.4 Allelic diversity at the S-locus within populations of B. insularis

The number of S-alleles observed within populations ranged from four in CA, the smallest population, to 18 for TG, the largest one. The intermediate sized populations (IZ and CG) showed 11 and 13 S-alleles, respectively (Fig. 2 and Fig. 4). Gene diversity at the S-locus, calculated as the expected heterozygosity (*He*), was low for CA (0.601), while it was higher and very similar among the three other populations, being 0.857, 0.863 and 0.897 for CG, IZ and TG respectively. As expected from theory (Schierup et al., 1997; Billiard et al., 2007), a recessive class II allele (H2002) reached the highest within-population frequency in most populations (CA, IZ and CG, Fig. 4). Intriguingly however, although this allele was also present in population TG, it was a dominant class I allele (H1028), that reached the highest frequency (Fig. 4).

**Figure 4.**
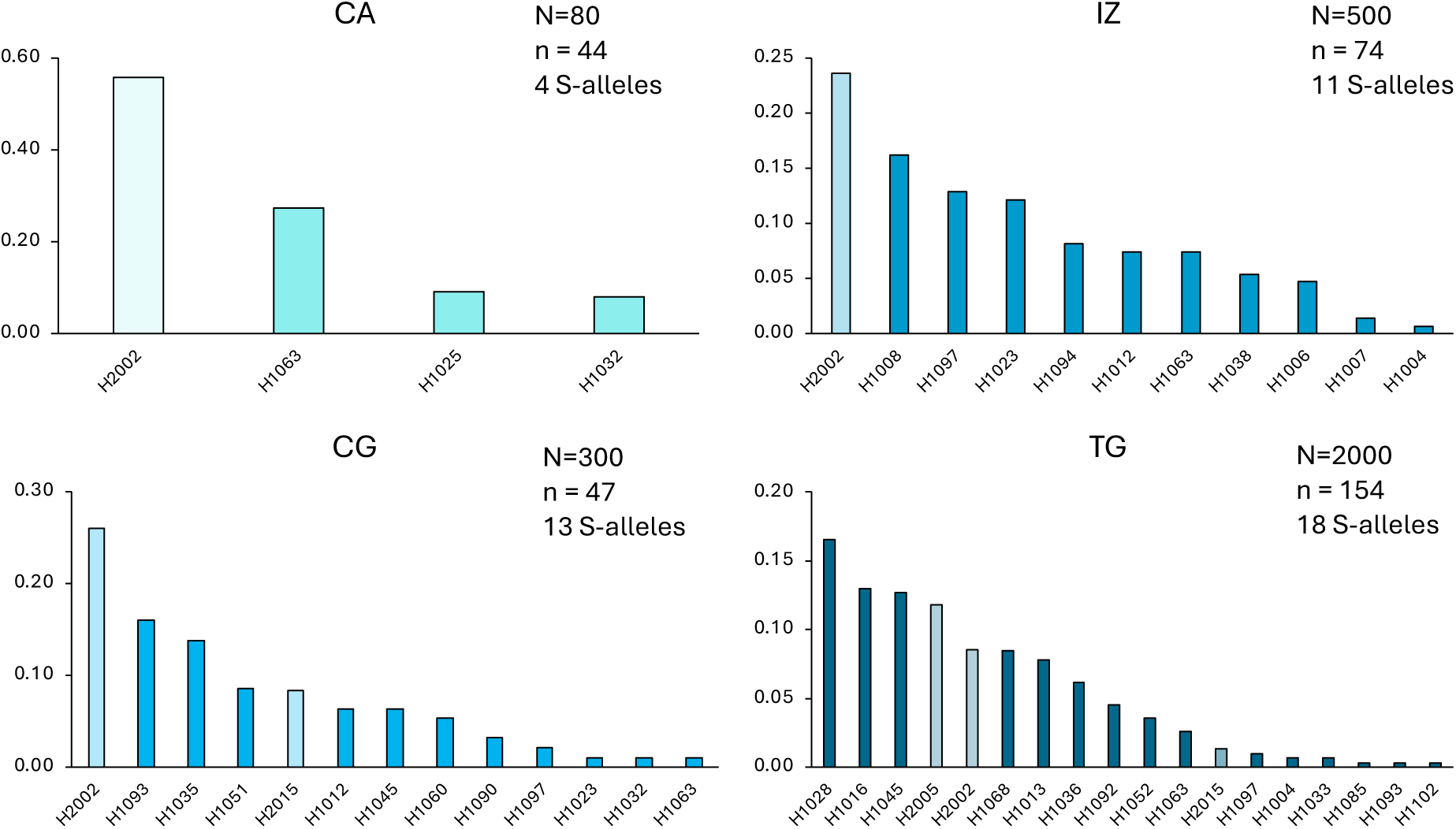
Frequency distribution of allelic frequencies within each population, with class I S-alleles in deeper color and class II S-alleles in lighter color. N is the estimated size of populations; n is the sample size.

Population genetic structure measured at the S-locus (*F*_st_ = 0.115) was lower than the value obtained by Glémin et al. (2005) for 11 microsatellite markers on the same populations (*F*_st_ =0.28, with confidence interval between 0.22 and 0.35). Remarkably, despite this low global *F*_ST_ value, 20 of the 31 S-alleles were found only in single populations, seven were found in two populations, one in three populations, whereas only two S-alleles, one class II and one class I, were common to all four populations (Fig.2).

The observed numbers of S-alleles per population were of the same order of magnitude as those expected for corresponding population sizes at a self-incompatibility locus under mutation-drift-selection equilibrium (Fig. 5). The fit was particularly good with expected values for a sporophytic SI system with dominance in pollen and codominance in stigma or for a gametophytic SI system, while the numbers of alleles expected for a sporophytic SI system with codominance in pollen and stigma were generally substantially higher than the observed values.

**Figure 5.**
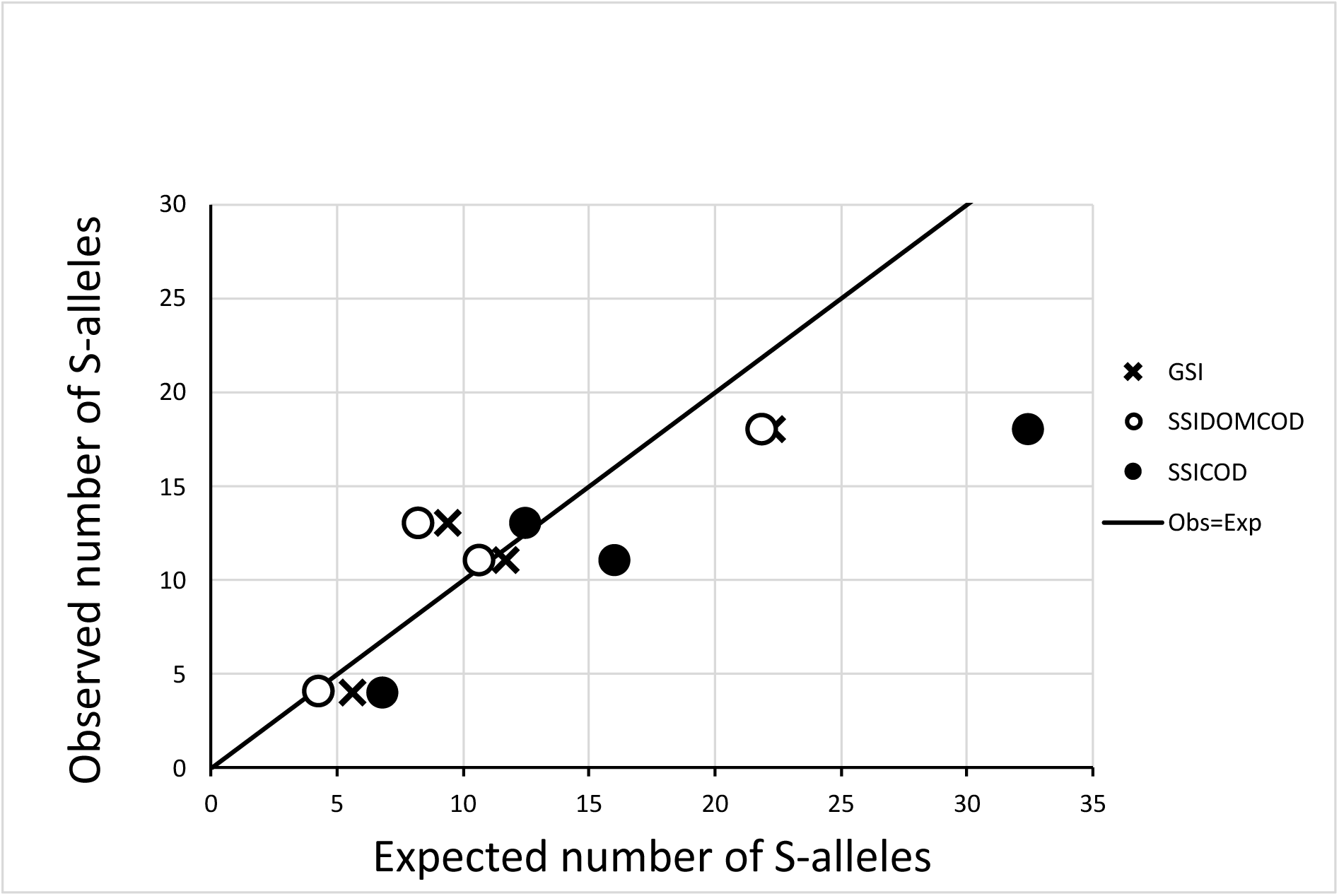
Comparison of observed and expected number of S-alleles in the four populations of *B. insularis*. Expected number of S-alleles are obtained either with analytical formula for Gametophytic SI (GSI), or with simulation under a sporophytic model with pollen limitation and either dominance in pollen and codominance in pistil (SSIDOMCOD) or codominance in both pollen and pistil (SSICOD).

### 3.5 Estimation of mate availability within populations of B. insularis

Based on individual genotypes at the S-locus, we estimated individual-based mate availability values from the female perspective, *i.e.* the proportion of individuals present in each population that are compatible as pollen donors with a given individual considered as a pollen receiver. Similarly to gene diversity values, but in a more pronounced way, average mate availability per population was found to be substantially lower in CA, the smallest population (mean=0.46), but was higher and rather homogeneous among the three other populations (mean for CG=0.72, IZ=0.68, TG=0.70; Fig. 6).

**Figure 6:**
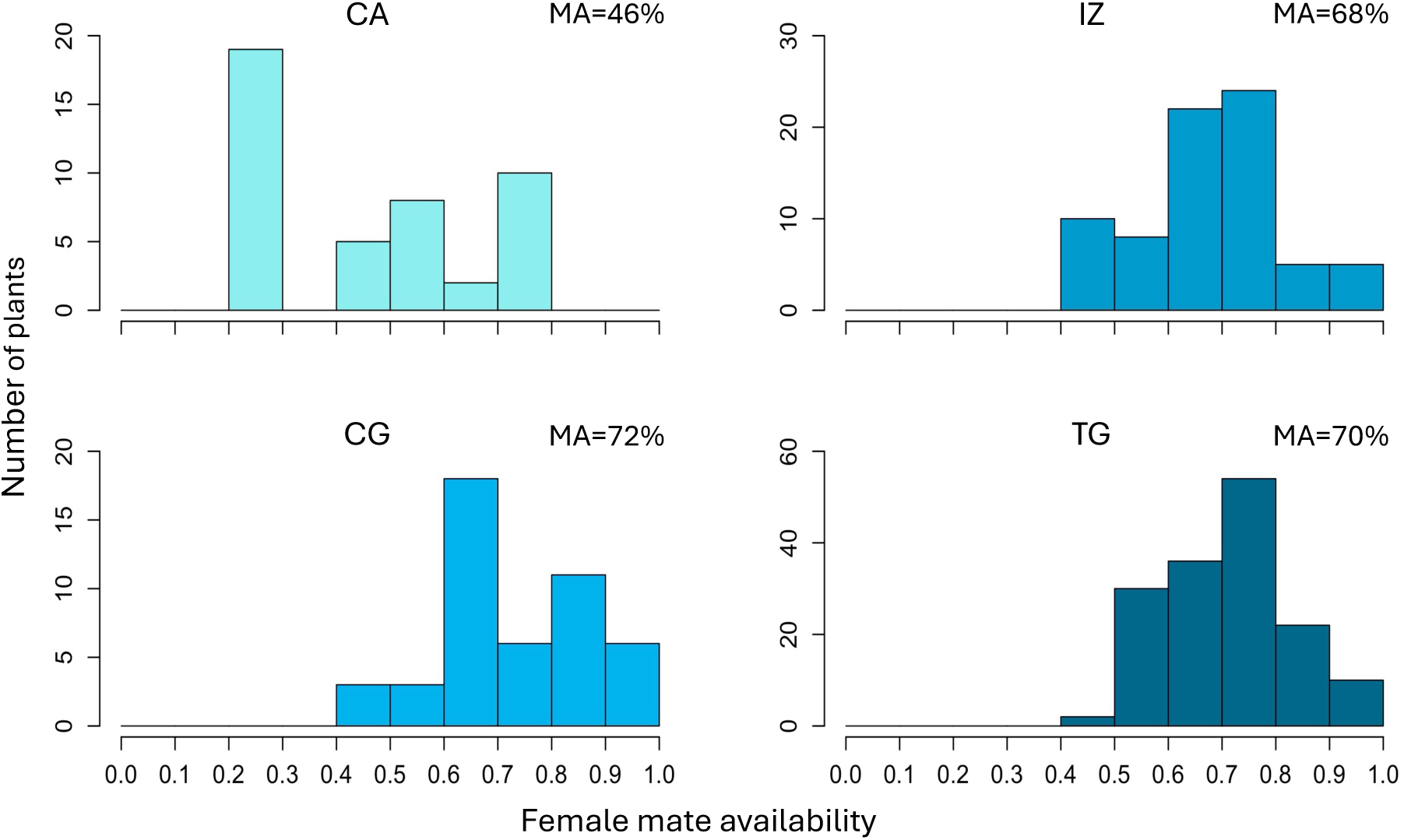
Distribution of individual mate availability within the four studied populations. MA is the population mean of individual female mate availability in percentage.

Strikingly, mate availability values showed strong variation among individuals, especially in the smallest population (standard deviation for CA=0.23, CG=0.14, IZ=0.12, TG=0.12). Indeed, in population CA, nineteen individuals showed a mate availability as low as 0.23 while the minimum individual value in other populations varied from 0.42 to 0.49 (Fig. 6).

## 4 DISCUSSION

### 4.1 Methodological approaches to genotype the S-locus in Brassica

With a few exceptions (e.g., Schierup et al., 2008, Glémin et al., 2005, Llaurens et al., 2008), most studies investigating the distribution of mate availability in natural populations of species with multiallelic SI have relied on phenotypic approaches to assess S-allele diversity (e.g. Byers, 1995; Campbell & Husband, 2007; Wagenius et al., 2007; Young & Pickup, 2010). However, these approaches are inherently limited to small sample sizes and preclude comprehensive estimation of S-allele diversity within and among populations.

The two NGS-based approaches that we tested in populations of *B. insularis* proved complementary. Together, they enabled us to obtain complete S-locus genotypes for the vast majority of individuals. The first approach (amplicon sequencing) was cost-effective but resulted in a substantial proportion (30%) of incomplete genotypes. These failures stemmed from a combination of complete amplification failure for four alleles, unreliable amplification for 11 alleles, and the inability to discriminate between two pairs of alleles due to high sequence identity in the targeted *SRK* region. Divergence at primer binding sites likely explain why some alleles were never detected, whereas inconsistent detection across individuals may reflect polymorphisms affecting amplification efficiency or competition among alleles. Methodological improvements could include the development of additional class I-specific primers targeting poorly amplified alleles.

The second approach, based on whole-genome resequencing, was more successful, yielding complete genotypes for all 246 individuals analyzed, except the two with low-quality DNA which could not be reanalyzed due to limited material. However, this approach is more costly. For data analysis, we used the bioinformatic pipeline NGSgenotyp, originally developed to genotype the S-locus in *Arabidopsis* species (Genete et al., 2020; Takou et al., 2021; Vekemans et al., 2026). Applying this pipeline to *Brassica* proved challenging for two main reasons. First, S-allele sequence divergence is lower in *Brassica* than in *Arabidopsis*, likely due to allelic re-diversification following the meso-polyploidization event at the origin of the *Brassica* tribe (Castric & Vekemans, 2007; Edh et al., 2009a). As a result, short reads from a given allele frequently map to closely related reference alleles—particularly within class II—leading to false positives. Second, the co-occurrence at the S-locus of two closely related genes (*SRK* and *SLG*), which undergo concerted evolution (Sato et al., 2002), results in cross-mapping of reads which biases mapping statistics (e.g. mismatch rates and depth of coverage). Together, these factors complicate the discrimination of closely related alleles and the identification of homozygous individuals. Modifying the pipeline by restricting analyses to reads with zero mismatches reduced ambiguity but did not fully resolve these issues. In contrast, using reference sequences of the pollen gene *SCR* instead of *SRK* allowed accurate genotyping with the NGSgenotyp pipeline. This strategy was successful and allowed to obtain full genotypes for 116 cultivated accessions of *B. oleracea*, as well as obtaining new SCR sequences for five S-alleles that were missing in GenBank. In *B. insularis* we identified 31 S-alleles, including four not previously reported in *B. oleracea* or *B. rapa*. This highlights the power of NGS-based methods, as novel alleles or missing sequences can be recovered through *de novo* assembly of filtered reads (Genete et al., 2020).

Although whole-genome resequencing is costly when targeting a single locus, it can leverage existing datasets generated for other purposes, such as those available for cultivated *Brassica* (Cheng et al., 2016; Zhang et al., 2018), and our successful analysis of 119 accessions of *B. oleracea* constitutes a proof of concept of this method. Moreover, in conservation studies, the genome-wide data produced can additionally provide robust estimates of genetic diversity and genetic load, offering valuable guidance for management strategies (Theissinger et al., 2023).

### 4.2 Species-level S-allele diversity in B. insularis as compared to other Brassica species

S-allele diversity has been studied in detail in *B. oleracea* and *B. rapa*, two commercially important species. Yamamoto et al. (2023) recently compiled sequences from 47 (44 of class I and 3 of class II) and 48 (44 of class I and 4 of class II) S-alleles in *B. oleracea* and *B. rapa*, respectively. In *B. cretica*, an endemic species from the Aegean archipelago phylogenetically close to *B. oleracea*, 21 S-alleles (18 of class I and 3 of class II) were observed in a total sample of only 40 individuals from four populations of Crete (Edh et al., 2009b). Here, using an extensive survey on 326 individuals from four populations of *B. insularis*, we observed 31 S-alleles (28 of class I and 3 of class II). The lower number of S-alleles observed in *B. cretica* as compared to the other Brassica species is probably associated with low sample size. In *B. insularis*, sample size is high locally, but the geographic range of the sampled populations (Corsica) is small as compared to the species distribution range (covering Corsica, Sardinia, Sicilia, Tunisia and Algeria). Moreover, we found that the proportion of allele sharing between populations was relatively low for class I alleles (68% of class I alleles were found in a single population), suggesting that many more class I alleles are probably occurring within the species.

Among the 31 S-alleles detected in *B. insularis*, 24 are shared with *B. oleracea* and three additional S-alleles are shared with *B.rapa* but not *B. oleracea,* based on combined evidence from *SCR* and *SRK* sequences (i.e. with nucleotide divergence lower than 2.5%). Functional maintenance of allelic specificity between trans-specific alleles of *B. oleracea* and *B. rapa* with high nucleotide similarity has been demonstrated for several allele pairs (Kimura et al., 2002; Sato et al., 2003). High degree of S-allele sharing among species is indeed expected due to the effect of strong negative frequency-dependent selection, which increases the coalescence times among allele lineages (i.e. it increases S-allele lifespan, Vekemans & Slatkin, 1994), and which also increases the chances of adaptive introgression of S-alleles among closely related species (Castric et al., 2008). It is particularly remarkable that the three class II alleles of *B. oleracea* are also found in *B. rapa* (Yamamoto et al., 2023), *B. cretica* (Edh et al. 2009b) and *B. insularis* (this study). This is in agreement with theoretical studies of sporophytic SI systems that predict higher allelic lifespan (Schierup et al., 1997; Vekemans et al., 1998) and wider among deme distribution in structured populations (Schierup et al., 2000) for recessive (i.e. class II) as compared to dominant (i.e. class I) alleles, particularly under situations of pollen limitation (Vekemans et al., 1998). Evidence for reduction in fruit set in populations of *B. insularis* suggests that pollen limitation is indeed occurring in this species (Glémin et al., 2008). Similarly, higher frequencies and wider among deme distribution of recessive as compared to dominant alleles was previously reported in populations of *Arabidopsis lyrata* (Schierup et al., 2008), which also expresses a sporophytic SI system, albeit with a larger number of dominance classes.

### 4.3 *S-allele* diversity within B. insularis populations and consequences for reproductive success

We found 4, 13, 11 and 18 S-alleles, respectively, for populations CA, CG, IZ and TG (in ranking order of estimated sizes). These numbers are quite similar to those estimated by Glémin et al (2005) based on biochemical markers, i.e. respectively 5, 12,11 and 15 for the same four populations, after correcting for the presence of null alleles. However, the number of S-alleles actually revealed species-wide was only 20, plus four alleles that the authors called ambiguous, as opposed to 31 in the present study. The order of magnitude of S-allele diversity within populations of *B. insularis*, when excluding population CA, is similar to those reported in other species with sporophytic SI, with values ranging between 11 and 14 in Icelandic (Schierup et al., 2008) and between 10 and 15 in continental (Takou et al., 2021) populations of *Arabidopsis lyrata*, and an estimate of 10 alleles in a French population of *A. halleri* (Llaurens et al., 2008).

The number of S-alleles found in each population of *B. insularis* increases with increasing population size, as expected from Wright’s finite population model of multiallelic SI (Wright, 1939). This was also observed by Young & Pickup (2010) in *Rutidosis leptorrhynchoides* (Asteraceae), with S-allele diversity, estimated from diallel crossing methods, ranging from 4 to 22 for population reproductive sizes ranging from 5 to 70.000. Dominance relationships between S-alleles in *Brassica* are intermediate between a purely dominant and a purely codominant model as dominance in pollen is observed between allelic classes but codominance is observed within classes (Hatakeyama et al., 1998; Yasuda et al., 2016). However, S-allele numbers were shown to fit better expectations based on simulations of a model with dominance in the pollen and codominance in the pistil (Fig. 5). Overall, the good fit with simulations considering single panmictic populations suggests that the Corsican populations of *B. insularis* are highly isolated, which is also in agreement with the relatively low fraction of shared alleles among populations. Data are also in accordance with the predictions that recessive class II alleles would be less numerous but more common than dominant class I alleles, and that class II alleles would reach a higher frequency when the class I alleles are less numerous (Uyenoyama, 2000).

Although the average mate availability was quite high in the three largest populations, around 70%, individual values were sometimes below 50%, suggesting potential reductions in reproductive output of some individuals. The situation is even worse in the smallest population which shows an average mate availability below 50%, with some individual values down to 23%, meaning that less than one fourth of the pollen sources are compatible with some individuals, these individuals representing more than 40% of the population (Fig. 6).

Mate availability values obtained here are higher than the estimations by Glémin et al (2008) based on their incomplete genotypes (e.g., 22% for CA and 59% for CG). Nevertheless, it is quite probable that at least in the smallest population, the fruit-set can be affected by the low mate availability. These results are comparable to those of Young and Pickup (2010) who used crosses to show that in seven populations of *R. leptorrhynchoides* mate availability was below 65% for the two populations of size less than 100 individuals. In Icelandic populations of *A. lyrata*, which are large continuous populations, mate availability was estimated to vary between 82 and 92% (Schierup et al., 2008).

As expected from theory (Schierup et al., 2000), populations are less structured at the S-locus (*F*_st_ = 0.115) than at neutral loci (*F*_st_ = 0.28, Glémin et al. 2005). However, the degree of allele sharing among populations was particularly low, with 20 alleles out of 31 found only in a single population, and only one class II and one class I alleles found in all four populations. This implies that creating some gene flow between populations would easily increase S-allele diversity and consequently enhance mate availability. Following a simulation study, Thrall et al. (2014) propose that adding new S-alleles in a population is profitable to its demography when the number of resident S-alleles is below 10-15. Nevertheless, whether this is recommended or not depends on the presence of local adaptation in the populations. Populations studied here live in quite different conditions (Noel et al., 2010), however the only quantitative genetics experiment available on this species showed a rather low structuring for quantitative traits (Petit et al., 2001) and the smallest population is in such a state that it may be worth the risk.

## AUTHORS CONTRIBUTIONS

Sandrine Maurice, Xavier Vekemans and Vincent Castric designed the study. Sandrine Maurice, Elodie Flaven, Agnès Mignot and Christophe Petit carried out the field work. Elodie Flaven performed the molecular work for the NGS amplicon-sequencing approach, Elodie Flaven and Sandrine Maurice analysed the genotypes using this method. Christelle Blassiau performed the molecular work for the whole genome resequencing approach and Mathieu Genete prepared the raw sequencing data for the bioinformatic analyses and adapted the NGSgenotyp pipeline. Mathieu Genete and Xavier Vekemans applied the pipeline to the resequencing data. Sandrine Maurice, Xavier Vekemans and Vincent Castric wrote the manuscript. All authors approved the final version.

## ACKNOWLEDGEMENTS

The authors thank Adeline Courseaux who made a preliminary test of the class I and class II primers designed in this study. Leaf materials have been collected with the contribution of the Conservatoire Botanique de Corse and the help of Eric Imbert.

## DATA AVAILABILITY STATEMENT

Supplementary material is available online. Reference sequences from *Brassica oleracea* and *B. rapa* used to analyze whole-genome resequencing data with the NGSgenotyp pipeline, as well as new *SRK* and *SCR* allele sequences obtained with the NGSgenotyp pipeline in *B. insularis* are available at https://doi.org/10.48579/PRO/EUP5ZX. The modified bioinformatic pipeline can be found at https://gitlab.univ-lille.fr/eep_lab/mathieu_genete/NGSgenotyp (https://doi.org/10.5281/zenodo.21506481). The code source for the individual-based simulation program used to investigate allelic diversity under SSI is available at https://gitlab.univ-lille.fr/eep_lab/xavier_vekemans/ssistruct. Short-read sequencing data obtained in this study are available at ENA under the project PRJEB122539.

