## Supplemental Figures and Tables for "Genotyping the self-incompatibility locus of wild and cultivated Brassica using NGS technologies: application to the evaluation of mate limitation in the endangered species *Brassica insularis*"

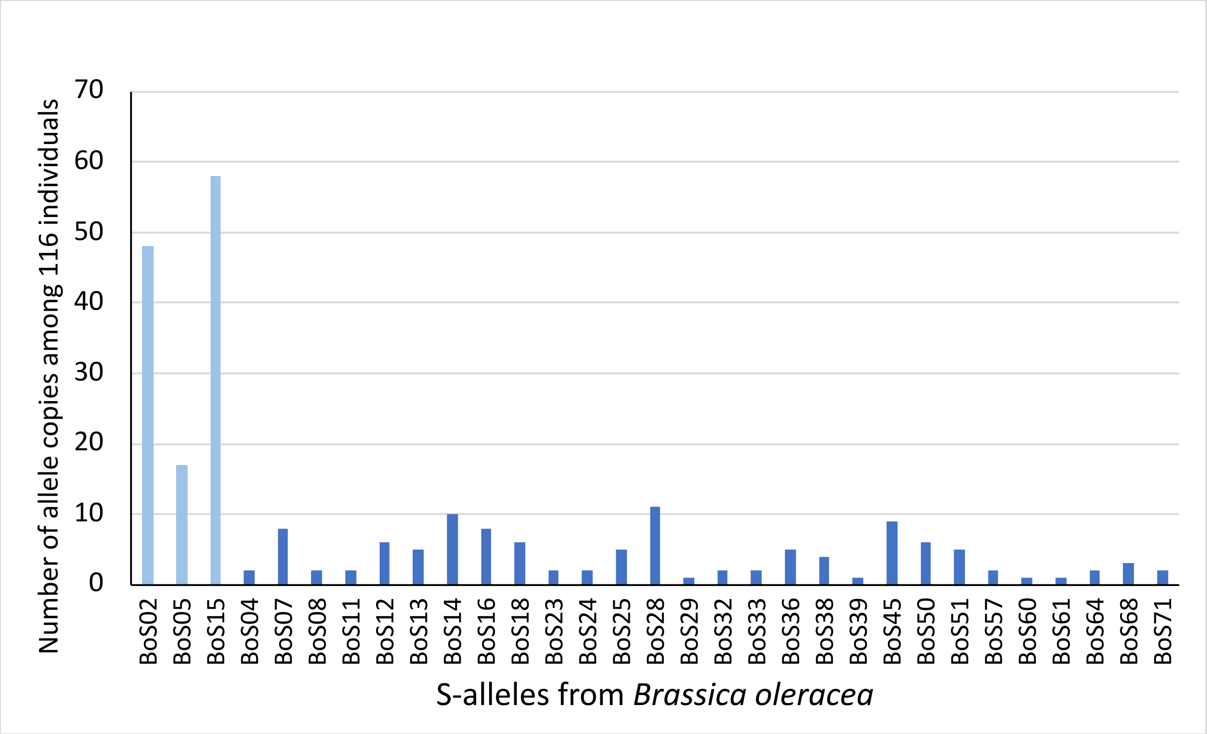


**Figure S1**. Distribution of S-allele frequencies among 116 accessions of *Brassica oleracea*, as determined using the NGSgenotyp pipeline on whole-genome resequencing data from Cheng et al., 2016). Class I alleles are represented with dark blue bars while class II alleles are represented by light blue bars.

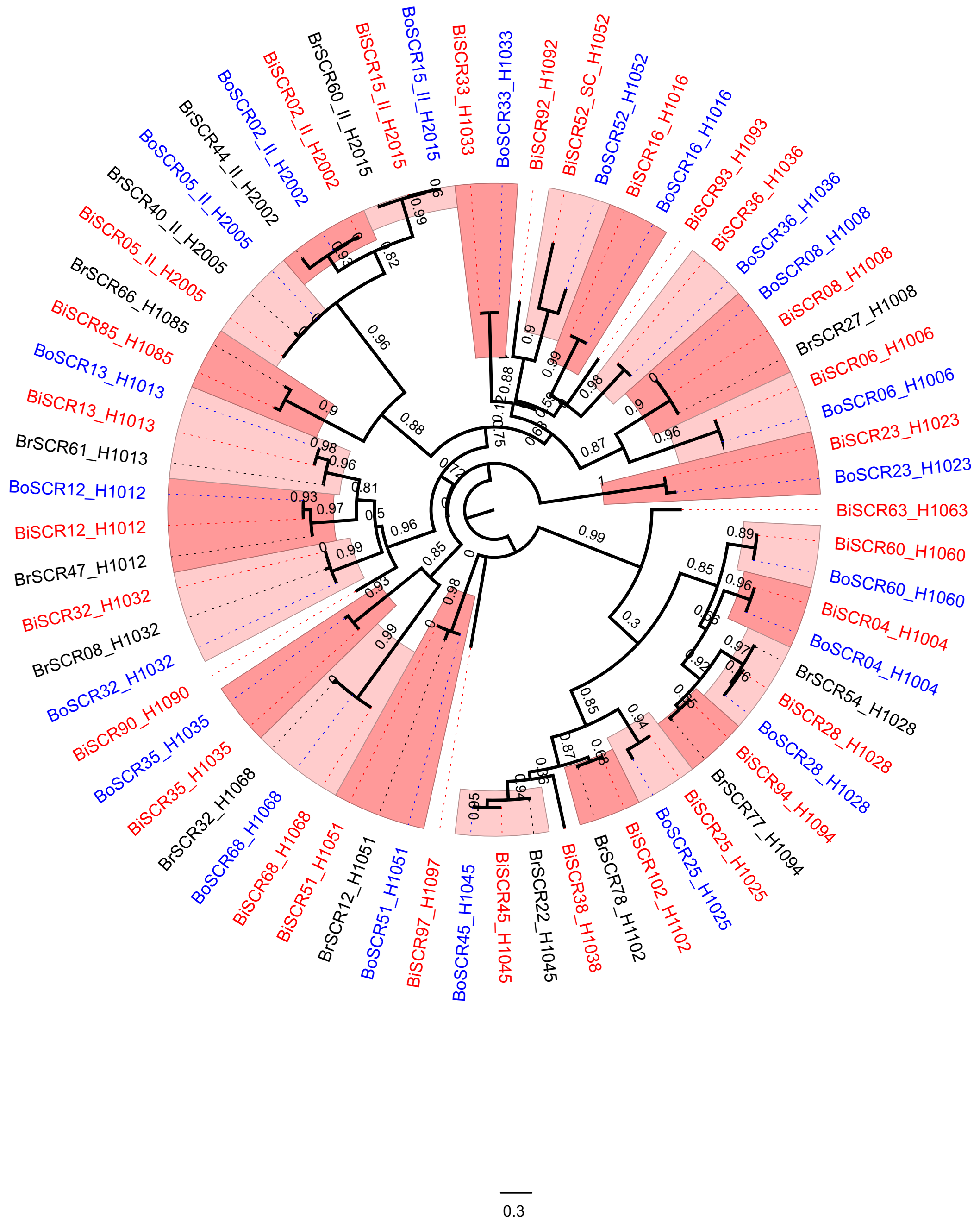


**Figure S2**. Phylogenies of *SCR* sequences from *Brassica insularis* (Bi, in red), *B. oleracea* (Bo, in blue) and *B. rapa* (Br, in black), as obtained using tree reconstruction based on maximum likelihood with PhyMl applied on amino acid sequences. Node support obtained from bootstraps is indicated. Groups of trans-specifically shared S-alleles are represented by alternate pink shades and labeled as HXYYY where X indicates whether the allele belongs to class I (1) or II (2), and YYY represents the label of the functional group.

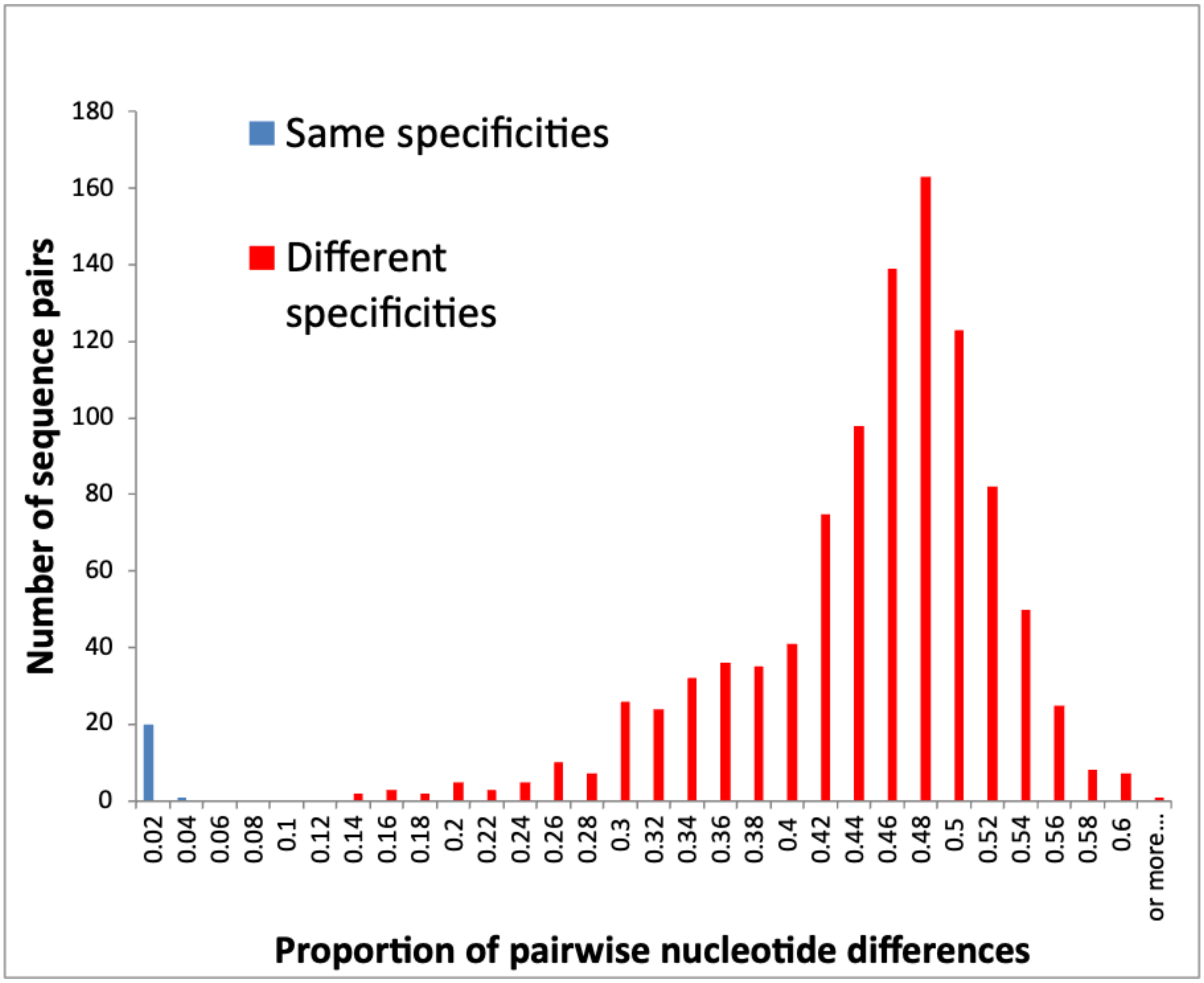


**Figure S3.** Distribution of pairwise nucleotide differences between *SCR* alleles from *Brassica insularis* and *B. oleracea* carrying the same specificity (blue bar) or different specificities (red bars)

**Table S1**. S-locus genotypes of 119 accessions from Brassica oleracea, sequenced by Cheng et al. (2016) and analyzed with our whole-genome resequencing approach using the NGSgenotyp pipeline.

| Sample | Ecotype | allele #1 | allele #2 | allele **#3** | SRA_accession |
|---|---|---|---|---|---|
| SamC_001 | Cabbage 1 | BoS15 | BoS15 | | SRR3202000 |
| SamC_002 | Cabbage 2 | BoS15 | BoS15 | | SRR3202020 |
| SamC_003 | Cabbage 3 | BoS02 | BoS02 | | SRR3202029 |
| SamC_004 | Cabbage 4 | BoS02 | BoS02 | | SRR3202031 |
| SamC_005 | Cabbage 5 | BoS15 | BoS15 | | SRR3202032 |
| SamC_006 | Cabbage 6 | BoS07 | BoS07 | | SRR3202033 |
| SamC_007 | Cabbage 7 | BoS50 | BoS50 | | SRR3202040 |
| SamC_008 | Cabbage 8 | BoS02 | BoS02 | | SRR3202041 |
| SamC_009 | Cabbage 9 | BoS14 | BoS14 | | SRR3202042 |
| SamC_010 | Cabbage 10 | BoS05 | BoS05 | | SRR3202043 |
| SamC_011 | Cabbage 11 | BoS02 | BoS02 | | SRR3202044 |
| SamC_012 | Cabbage 12 | BoS45 | BoS45 | | SRR3202802 |
| SamC_013 | Cabbage 13 | BoS68 | BoS02 | | SRR3202803 |
| SamC_014 | Cabbage 14 | BoS15 | BoS15 | | SRR3202805 |
| SamC_015 | Cabbage 15 | BoS02 | BoS02 | | SRR3202806 |
| SamC_016 | Cabbage 16 | BoS28 | BoS28 | | SRR3202807 |
| SamC_017 | Cabbage 17 | BoS07 | BoS07 | | SRR3202808 |
| SamC_018 | Cabbage 18 | BoS05 | BoS05 | | SRR3202809 |
| SamC_019 | Cabbage 19 | BoS57 | BoS57 | | SRR3202810 |
| SamC_020 | Cabbage 20 | BoS45 | BoS45 | | SRR3202811 |
| SamC_021 | Cabbage 21 | BoS02 | BoS02 | | SRR3202820 |
| SamC_022 | Cabbage 22 | BoS05 | BoS05 | | SRR3202827 |
| SamC_023 | Cabbage 23 | BoS28 | BoS28 | | SRR3202833 |
| SamC_024 | Cabbage 24 | BoS02 | BoS02 | | SRR3202834 |
| SamC_025 | Cabbage 25 | BoS51 | BoS51 | | SRR3202835 |
| SamC_026 | Cabbage 26 | BoS28 | BoS28 | | SRR3202836 |
| SamC_027 | Cabbage 27 | BoS33 | BoS33 | | SRR3202837 |
| SamC_028 | Cabbage 28 | BoS36 | BoS36 | | SRR3202838 |
| SamC_029 | Cabbage 29 | BoS14 | BoS50 | | SRR3202868 |
| SamC_030 | Pointed Cabbage 1 | BoS02 | BoS02 | | SRR3202869 |
| SamC_031 | Pointed Cabbage 2 | BoS05 | BoS05 | | SRR3202870 |
| SamC_032 | Pointed Cabbage 3 | BoS25 | BoS25 | | SRR3202873 |
| SamC_033 | White Cabbage 1 | BoS15 | BoS15 | | SRR3202875 |
| SamC_034 | White Cabbage 2 | BoS14 | BoS14 | | SRR3202876 |
| SamC_035 | White Cabbage 3 | BoS28 | BoS28 | | SRR3202877 |

| | | | | | |
|---|---|---|---|---|---|
| SamC_036 | White Cabbage 4 | BoS07 | BoS07 | | SRR3202881 |
| SamC_037 | White Cabbage 5 | BoS50 | BoS50 | | SRR3202882 |
| SamC_038 | White Cabbage 6 | BoS14 | BoS50 | | SRR3202883 |
| SamC_039 | White Cabbage 7 | BoS07 | BoS07 | | SRR3202885 |
| SamC_040 | White Cabbage 8 | BoS38 | BoS38 | | SRR3202912 |
| SamC_041 | White Cabbage 9 | BoS15 | BoS15 | | SRR3203045 |
| SamC_042 | White Cabbage 10 | BoS15 | BoS15 | | SRR3203047 |
| SamC_043 | White Cabbage 11 | BoS05 | BoS05 | | SRR3203048 |
| SamC_044 | Cabbage 30 | BoS51 | BoS02 | | SRR3203049 |
| SamC_045 | Cabbage 31 | BoS02 | BoS05 | | SRR3203050 |
| SamC_046 | Kohlrabi 1 | BoS12 | BoS12 | | SRR3203051 |
| SamC_047 | Kohlrabi 2 | BoS04 | BoS04 | | SRR3203052 |
| SamC_048 | Kohlrabi 3 | BoS15 | BoS15 | | SRR3203053 |
| SamC_049 | Kohlrabi 4 | BoS24 | BoS24 | | SRR3203054 |
| SamC_050 | Kohlrabi 5 | BoS64 | BoS64 | | SRR3203055 |
| SamC_051 | Kohlrabi 6 | missing | missing | | SRR3203056 |
| SamC_052 | Kohlrabi 7 | BoS61 | BoS15 | | SRR3203072 |
| SamC_053 | Kohlrabi 8 | BoS28 | BoS15 | | SRR3203074 |
| SamC_054 | Kohlrabi 9 | BoS16 | BoS16 | | SRR3203075 |
| SamC_055 | Kohlrabi 10 | BoS15 | BoS15 | | SRR3203076 |
| SamC_056 | Kohlrabi 11 | BoS15 | BoS15 | | SRR3203077 |
| SamC_057 | Kohlrabi 12 | BoS02 | BoS02 | | SRR3203079 |
| SamC_058 | Kohlrabi 13 | BoS16 | BoS16 | | SRR3203080 |
| SamC_059 | Kohlrabi 14 | BoS16 | BoS16 | | SRR3203082 |
| SamC_060 | Kohlrabi 15 | BoS05 | BoS05 | | SRR3203083 |
| SamC_061 | Kohlrabi 16 | BoS11 | BoS11 | | SRR3203084 |
| SamC_062 | Kohlrabi 17 | BoS12 | BoS12 | | SRR3203086 |
| SamC_063 | Kohlrabi 18 | BoS68 | BoS68 | | SRR3203087 |
| SamC_064 | Kohlrabi 19 | BoS71 | BoS71 | | SRR3203089 |
| SamC_065 | Cauliflower 1 | BoS15 | BoS15 | | SRR3203096 |
| SamC_066 | Cauliflower 2 | BoS15 | BoS15 | | SRR3203104 |
| SamC_067 | Cauliflower 3 | BoS36 | BoS36 | | SRR3203105 |
| SamC_068 | Cauliflower 4 | BoS15 | BoS15 | | SRR3203106 |
| SamC_069 | Cauliflower 5 | BoS15 | BoS15 | | SRR3203107 |
| SamC_070 | Cauliflower 6 | BoS15 | BoS15 | | SRR3203108 |
| SamC_071 | Cauliflower 7 | BoS14 | BoS14 | | SRR3203109 |
| SamC_072 | Cauliflower 8 | BoS02 | BoS02 | | SRR3203115 |
| SamC_073 | Cauliflower 9 | BoS14 | BoS14 | | SRR3203119 |
| SamC_074 | Cauliflower 10 | BoS08 | BoS08 | | SRR3203120 |

| | | | | | |
|---|---|---|---|---|---|
| SamC_075 | Cauliflower 11 | BoS15 | BoS15 | | SRR3203127 |
| SamC_076 | Cauliflower 12 | BoS23 | BoS23 | | SRR3203131 |
| SamC_077 | Cauliflower 13 | BoS05 | BoS05 | | SRR3203136 |
| SamC_078 | Cauliflower 14 | BoS15 | BoS15 | | SRR3203140 |
| SamC_079 | Cauliflower 15 | BoS15 | BoS15 | | SRR3203144 |
| SamC_080 | Cauliflower 16 | BoS25 | BoS45 | | SRR3203147 |
| SamC_081 | Cauliflower 17 | BoS16 | BoS16 | | SRR3203151 |
| SamC_082 | Cauliflower 18 | BoS02 | BoS02 | | SRR3203155 |
| SamC_083 | Cauliflower 19 | BoS45 | BoS45 | | SRR3203157 |
| SamC_084 | Cauliflower 20 | BoS12 | BoS12 | | SRR3203160 |
| SamC_085 | Broccoli 1 | BoS02 | BoS02 | | SRR3203163 |
| SamC_086 | Broccoli 2 | BoS15 | BoS15 | | SRR3203164 |
| SamC_087 | Broccoli 3 | BoS13 | BoS13 | | SRR3203165 |
| SamC_088 | Broccoli 4 | BoS15 | BoS15 | | SRR3203166 |
| SamC_089 | Broccoli 5 | BoS15 | BoS15 | | SRR3203169 |
| SamC_090 | Broccoli 6 | BoS18 | BoS18 | | SRR3203172 |
| SamC_091 | Broccoli 7 | BoS18 | BoS18 | | SRR3203176 |
| SamC_092 | Broccoli 8 | BoS02 | BoS02 | | SRR3203181 |
| SamC_093 | Broccoli 9 | BoS15 | BoS15 | | SRR3203187 |
| SamC_094 | Broccoli 10 | BoS02 | BoS02 | | SRR3203201 |
| SamC_095 | Broccoli 11 | BoS36 | BoS02 | BoS15 | SRR3203204 |
| SamC_096 | Broccoli 12 | BoS51 | BoS39 | BoS02 | SRR3203206 |
| SamC_097 | Broccoli 13 | BoS15 | BoS15 | | SRR3203212 |
| SamC_098 | Broccoli 14 | BoS15 | BoS15 | | SRR3203219 |
| SamC_099 | Broccoli 15 | BoS02 | BoS02 | | SRR3203228 |
| SamC_100 | Broccoli 16 | BoS02 | BoS02 | | SRR3203237 |
| SamC_101 | Broccoli 17 | BoS15 | BoS15 | | SRR3203244 |
| SamC_102 | Broccoli 18 | BoS02 | BoS02 | | SRR3203255 |
| SamC_103 | Broccoli 19 | BoS13 | BoS13 | | SRR3203257 |
| SamC_104 | Broccoli 20 | BoS18 | BoS18 | | SRR3203258 |
| SamC_105 | Broccoli 21 | BoS15 | BoS15 | | SRR3203259 |
| SamC_106 | Broccoli 22 | BoS02 | BoS02 | | SRR3203260 |
| SamC_107 | Broccoli 23 | BoS15 | BoS15 | | SRR3203261 |
| SamC_108 | Chinese kale 1 | BoS05 | BoS05 | | SRR3203263 |
| SamC_109 | Chinese kale 2 | BoS28 | BoS28 | | SRR3203264 |
| SamC_110 | Chinese kale 3 | BoS25 | BoS25 | | SRR3203269 |
| SamC_111 | Chinese kale 4 | BoS02 | BoS02 | | SRR3203270 |
| SamC_112 | Brussels sprouts 1 | BoS45 | BoS45 | | SRR3203273 |
| SamC_113 | Brussels sprouts 2 | BoS60 | BoS02 | | SRR3203278 |

| | | | | | |
|---|---|---|---|---|---|
| SamC_114 | Kale 1 | BoS02 | BoS02 | | SRR3203281 |
| SamC_115 | Kale 2 | BoS29 | BoS02 | | SRR3203286 |
| SamC_116 | Curly Kale 1 | BoS38 | BoS38 | | SRR3203293 |
| SamC_117 | Curly Kale 2 | BoS32 | BoS32 | | SRR3203366 |
| SamC_118 | Wild 1 | BoS51 | BoS15 | | SRR3203367 |
| SamC_119 | Wild 2 | BoS13 | BoS02 | | SRR3203368 |

**Table S2.** List of the 31 S-alleles identified in Brassica oleracea accessions. The accessions used to extract the five new SCR exon 2 sequences using the NGSgenotyp pipeline are indicated. The Allele ID corresponds to a new identification scheme based on nucleotide divergence among SRK and SCR sequences from different Brassica species, in the form HXYYY, with X corresponding to the dominance class of the allele (I = 1; II = 2), and YYY corresponding to its functional specificity shared among species (ranging from 001 up to a limit of 999).

| Allele Class | Allele ID (functional group) | *B. oleracea* specific allele ID | ID of new *SCR* sequences | Accesion used to extract *SCR* sequence |
|---|---|---|---|---|
| I | H1004 | BoS04 | | |
| I | H1007 | BoS07 | | |
| I | H1008 | BoS08 | | |
| I | H1011 | BoS11 | | |
| I | H1012 | BoS12 | | |
| I | H1013 | BoS13 | | |
| I | H1014 | BoS14 | | |
| I | H1016 | BoS16 | | |
| I | H1018 | BoS18 | | |
| I | H1023 | BoS23 | BoSCR23 | SamC_076 |
| I | H1024 | BoS24 | | |
| I | H1025 | BoS25 | | |
| I | H1028 | BoS28 | | |
| I | H1029 | BoS29 | | |
| I | H1032 | BoS32 | | |
| I | H1033 | BoS33 | | |
| I | H1036 | BoS36 | BoSCR36 | SamC_067 |
| I | H1038 | BoS38 | BoSCR38 | SamC_040 |
| I | H1039 | BoS39 | | |
| I | H1045 | BoS45 | | |
| I | H1050 | BoS50 | BoSCR50 | SamC_037 |
| I | H1051 | BoS51 | | |
| I | H1057 | BoS57 | | |
| I | H1060 | BoS60 | BoSCR60 | SamC_113 |
| I | H1061 | BoS61 | | |
| I | H1064 | BoS64 | | |
| I | H1068 | BoS68 | | |
| I | H1071 | BoS71 | | |
| II | H2002 | BoS02 | | |
| II | H2005 | BoS05 | | |
| II | H2015 | BoS15 | | |

**Table S3.** List of the 31 S-alleles identified in *Brassica insularis*, with correspondence to known alleles in *B. oleracea* and *B. rapa*, based on sequence similarity and phylogenetic relationships. The individuals used to extract the *SRK* S-domain sequence (exon 1) and the *SCR* exon 2 of each *B. insularis* allele using the NGSgenotyp pipeline are indicated. The Allele ID corresponds to a new identification scheme based on nucleotide divergence among *SRK* and *SCR* sequences from different Brassica species, in the form HXYYY, with X corresponding to the dominance class of the allele (I = 1; II = 2), and YYY corresponding to its functional specificity shared among species (ranging from 001 up to a limit of 999).

| Allele Class | Allele ID (functional group) | *B. insularis* specific allele ID | *B. oleracea* homolog | *B. rapa* homolog | ID of new *SRK* sequences | Individual used to extract *SRK* sequence | ID of new *SCR* sequences | Individual used to extract *SCR* sequence |
|---|---|---|---|---|---|---|---|---|
| I | H1004 | BiS04 | BoS04 | | BiSRK04 | TG301_Contig_N9 | BiSCR04 | TG301_contig=6011 |
| I | H1006 | BiS06 | BoS06 | | BiSRK06 | IZ184_Contig_N6 | BiSCR06 | IZ156_Contig=8219 |
| I | H1007 | BiS07 | BoS07 | BrS46 | BiSRK07 | IZ114_Contig_N28 | BiSCR07 | IZ112 |
| I | H1008 | BiS08 | BoS08 | BrS27 | BiSRK08 | IZ03_Contig_N4 | BiSCR08 | IZ147_contig=1965 |
| I | H1012 | BiS12 | BoS12 | BrS47 | BiSRK12 | CG94_Contig_N5 | BiSCR12 | CG94_Contig=N2095 |
| I | H1013 | BiS13 | BoS13 | BrS61 | BiSRK13 | TG186_Contig_N2 | BiSCR13 | TG192_contig=131 |
| I | H1016 | BiS16 | BoS16 | | BiSRK16 | TG103_Contig_N1 | BiSCR16 | TG107_Contig=2159 |
| I | H1023 | BiS23 | BoS23 | | BiSRK23 | IZ168_Contig_N3 | BiSCR23 | IZ181_Contig=8755 |
| I | H1025 | BiS25 | BoS25 | | BiSRK25 | CA88_Contig_N13 | BiSCR25 | CA88_Contig=7971 |
| I | H1028 | BiS28 | BoS28 | BrS54 | BiSRK28 | TG106_Contig_N2 | BiSCR28 | TG103_Contig=1512 |
| I | H1032 | BiS32 | BoS32 | BrS08 | BiSRK32 | CA11_Contig_N6 | BiSCR32 | CA52_Contig=N2710 |
| I | H1033 | BiS33 | BoS33 | | BiSRK33 | TG111_Contig_N10 | BiSCR33 | TG111_Contig=3402 |
| I | H1035 | BiS35 | BoS35 | | BiSRK35 | CG154_Contig_N4 | BiSCR35 | CG94_Contig=N1737 |
| I | H1036 | BiS36 | BoS36 | | BiSRK36 | TG121_Contig_N12 | BiSCR36 | TG127_contig=1451 |
| I | H1038 | BiS38 | BoS38 | | BiSRK38 | IZ148_Contig_N13 | BiSCR38 | IZ100_Contig=3125 |
| I | H1045 | BiS45 | BoS45 | BrS22 | BiSRK45 | TG320_Contig_N3 | BiSCR45 | TG86_Contig=2402 |
| I | H1051 | BiS51 | BoS51 | BrS12 | BiSRK51 | CG89_Contig_N8 | BiSCR51 | CG154_contig=1137 |
| I | H1052 | BiS52 | BoS52 | | BiSRK52 | TG112_Contig_N6 | BiSCR52 | TG121_contig=4970 |
| I | H1060 | BiS60 | BoS60 | | BiSRK60 | CG77_Contig_N14 | BiSCR60 | CG114_contig=4293 |
| I | H1063 | BiS63 | BoS63 | | BiSRK63 | TG281_Contig_N3 | BiSCR63 | TG93_contig=2326 |
| I | H1068 | BiS68 | BoS68 | BrS32 | BiSRK68 | TG124_Contig_N1 | BiSCR68 | TG124_Contig=1346 |
| I | H1085 | BiS85 | | BrS66 | BiSRK85 | TG135_Contig_N8 | BiSCR85 | TG135_contig=1913 |
| I | H1090 | BiS90 | | | BiSRK90 | CG119_Contig_N4 | BiSCR90 | CG108_Contig=6817 |
| I | H1092 | BiS92 | | | BiSRK92 | TG104_Contig_N4 | BiSCR92 | TG86_contig=407 |
| I | H1093 | BiS93 | | | BiSRK93 | CG152_Contig_N4 | BiSCR93 | CG134_contig=17176 |
| I | H1094 | BiS94 | | BrS77 | BiSRK94 | IZ125_Contig_N6 | BiSCR94 | IZ104_contig=4131 |
| I | H1097 | BiS97 | | | BiSRK97 | CG93_Contig_N1 | BiSCR97 | IZ183_Contig=9268 |
| I | H1102 | BiS102 | | BrS78 | BiSRK98 | CG159_Contig_N44 | BiSCR102 | TG108_contig=1328 |
| II | H2002 | BiS02 | BoS02 | BrS44 | BiSRK02 | IZ03_Contig_N5 | BiSCR02 | CA25_Contig=N84 |
| II | H2005 | BiS05 | BoS05 | BrS40 | BiSRK05 | TG253_Contig_N2 | BiSCR05 | TG230_contig=1854 |
| II | H2015 | BiS15 | BoS15 | BrS60 | BiSRK15 | CG146_Contig_N8 | BiSCR15 | CG122_contig=488 |

**Table S4.** S-locus genotypes of 319 individuals from *Brassica insularis*, as obtained by combining an amplicon sequencing approach and a whole-genome resequencing approach. For the latter, we indicate the SRA accession ID corresponding to the sequencing data obtained in this study. S-allele sequences are identified according to a new *Brassica* functional sequence group nomenclature, as determined based on sequence similarity with alleles from the related *Brassica oleracea* and *B. rapa* (see Table S3 for correspondence with *B. insularis*-specific S-allele IDs).

| Sample | Population | allele #1 | allele #2 | SRA_accession |
|---|---|---|---|---|
| CA06 | Punta Calcina (CA) | H2002 | H2002 | ERR17631665 |
| CA09 | Punta Calcina (CA) | H1025 | H2002 | |
| CA10 | Punta Calcina (CA) | H1025 | H2002 | |
| CA11 | Punta Calcina (CA) | H1032 | H1063 | ERR17631696 |
| CA12 | Punta Calcina (CA) | H1032 | H1063 | |
| CA15 | Punta Calcina (CA) | H2002 | H2002 | ERR17631652 |
| CA25 | Punta Calcina (CA) | H2002 | H2002 | ERR17631678 |
| CA28 | Punta Calcina (CA) | H1063 | H2002 | |
| CA30 | Punta Calcina (CA) | H1063 | H2002 | ERR17631779 |
| CA31 | Punta Calcina (CA) | H1063 | H2002 | |
| CA38 | Punta Calcina (CA) | H1063 | H2002 | |
| CA42 | Punta Calcina (CA) | H1032 | H1063 | ERR17631677 |
| CA45 | Punta Calcina (CA) | H1032 | H1063 | ERR17631569 |
| CA52 | Punta Calcina (CA) | H1032 | H1063 | ERR17631734 |
| CA58 | Punta Calcina (CA) | H2002 | H2002 | ERR17631657 |
| CA60 | Punta Calcina (CA) | H1063 | H2002 | |
| CA61 | Punta Calcina (CA) | H2002 | H2002 | ERR17631712 |
| CA62 | Punta Calcina (CA) | H1032 | H2002 | |
| CA66 | Punta Calcina (CA) | H2002 | H2002 | |
| CA67 | Punta Calcina (CA) | H1063 | H2002 | ERR17631675 |
| CA68 | Punta Calcina (CA) | H2002 | H2002 | ERR17631669 |
| CA72 | Punta Calcina (CA) | H1063 | H2002 | ERR17631727 |
| CA74 | Punta Calcina (CA) | H1063 | H2002 | |
| CA75 | Punta Calcina (CA) | H1063 | H2002 | |
| CA76 | Punta Calcina (CA) | H1025 | H2002 | |
| CA77 | Punta Calcina (CA) | H1063 | H2002 | |
| CA78 | Punta Calcina (CA) | H1063 | H2002 | |
| CA80 | Punta Calcina (CA) | H2002 | H2002 | ERR17631556 |

| | | | | |
|---|---|---|---|---|
| CA82 | Punta Calcina (CA) | H1032 | H2002 | |
| CA83 | Punta Calcina (CA) | H1063 | H2002 | |
| CA84 | Punta Calcina (CA) | H1063 | H2002 | ERR17631715 |
| CA85 | Punta Calcina (CA) | H1063 | H2002 | |
| CA86 | Punta Calcina (CA) | H1063 | H2002 | |
| CA87 | Punta Calcina (CA) | H1025 | H2002 | |
| CA88 | Punta Calcina (CA) | H1025 | H2002 | |
| CA89 | Punta Calcina (CA) | H2002 | H2002 | ERR17631539 |
| CA90 | Punta Calcina (CA) | H2002 | H2002 | ERR17631725 |
| CA91 | Punta Calcina (CA) | H1025 | H2002 | |
| CA92 | Punta Calcina (CA) | H1063 | H2002 | |
| CA109 | Punta Calcina (CA) | H1063 | H2002 | |
| CA110 | Punta Calcina (CA) | H1063 | H2002 | ERR17631586 |
| CA115 | Punta Calcina (CA) | H1063 | H2002 | ERR17631648 |
| CA117 | Punta Calcina (CA) | H1025 | H2002 | ERR17631594 |
| CA118 | Punta Calcina (CA) | H1025 | H2002 | ERR17631776 |
| CG36 | Punta Corbaghiola (CG) | H1060 | H2002 | |
| CG37 | Punta Corbaghiola (CG) | H1060 | H2002 | ERR17631719 |
| CG62 | Punta Corbaghiola (CG) | H1012 | H1060 | ERR17631702 |
| CG77 | Punta Corbaghiola (CG) | H1032 | H1060 | ERR17631537 |
| CG88 | Punta Corbaghiola (CG) | H1045 | H1035 | |
| CG89 | Punta Corbaghiola (CG) | H1051 | H1035 | ERR17631574 |
| CG90 | Punta Corbaghiola (CG) | H1035 | H1093 | |
| CG92 | Punta Corbaghiola (CG) | H1051 | H1035 | |
| CG93 | Punta Corbaghiola (CG) | H1097 | H1093 | ERR17631767 |
| CG94 | Punta Corbaghiola (CG) | H1012 | H1035 | ERR17631767 |
| CG95 | Punta Corbaghiola (CG) | H1012 | H1035 | ERR17631714 |
| CG96 | Punta Corbaghiola (CG) | H1035 | H1093 | ERR17631706 |
| CG99 | Punta Corbaghiola (CG) | H1035 | H2002 | ERR17631689 |
| CG100 | Punta Corbaghiola (CG) | H1093 | H2002 | ERR17631769 |
| CG101 | Punta Corbaghiola (CG) | H1093 | H2002 | |
| CG104 | Punta Corbaghiola (CG) | H1093 | H2002 | |
| CG105 | Punta Corbaghiola (CG) | H1035 | H1093 | |
| CG106 | Punta Corbaghiola (CG) | H2002 | H2015 | ERR17631559 |
| CG108 | Punta Corbaghiola (CG) | H1090 | H2015 | ERR17631647 |
| CG109 | Punta Corbaghiola (CG) | H1093 | H2002 | |
| CG112 | Punta Corbaghiola (CG) | H1097 | H1012 | ERR17631711 |
| CG114 | Punta Corbaghiola (CG) | H1060 | H2002 | ERR17631713 |
| CG118 | Punta Corbaghiola (CG) | H1045 | H2015 | |

| | | | | |
|---|---|---|---|---|
| CG119 | Punta Corbaghiola (CG) | H1090 | H2002 | ERR17631738 |
| CG120 | Punta Corbaghiola (CG) | H1051 | H2002 | |
| CG122 | Punta Corbaghiola (CG) | H1045 | H2015 | ERR17631716 |
| CG126 | Punta Corbaghiola (CG) | H1051 | H2002 | |
| CG127 | Punta Corbaghiola (CG) | H1051 | H2002 | |
| CG129 | Punta Corbaghiola (CG) | H1063 | H1090 | ERR17631664 |
| CG134 | Punta Corbaghiola (CG) | H1093 | H2002 | ERR17631737 |
| CG140 | Punta Corbaghiola (CG) | H1051 | H1093 | ERR17631701 |
| CG142 | Punta Corbaghiola (CG) | H1035 | H2002 | ERR17631567 |
| CG143 | Punta Corbaghiola (CG) | H1051 | H2002 | |
| CG144 | Punta Corbaghiola (CG) | H1045 | H2002 | |
| CG146 | Punta Corbaghiola (CG) | H1045 | H2015 | ERR17631728 |
| CG147 | Punta Corbaghiola (CG) | H1093 | H2002 | ERR17631621 |
| CG148 | Punta Corbaghiola (CG) | H1035 | H2002 | ERR17631544 |
| CG149 | Punta Corbaghiola (CG) | H1093 | H1012 | ERR17631668 |
| CG150 | Punta Corbaghiola (CG) | H2002 | H2015 | ERR17631630 |
| CG151 | Punta Corbaghiola (CG) | H1045 | H1093 | ERR17631761 |
| CG152 | Punta Corbaghiola (CG) | H1093 | H2002 | ERR17631582 |
| CG153 | Punta Corbaghiola (CG) | H1035 | H2015 | ERR17631751 |
| CG154 | Punta Corbaghiola (CG) | H1035 | H1051 | ERR17631592 |
| CG158 | Punta Corbaghiola (CG) | H2002 | H2015 | ERR17631786 |
| CG160 | Punta Corbaghiola (CG) | H1093 | H2002 | ERR17631670 |
| CG161 | Punta Corbaghiola (CG) | H1012 | H2002 | ERR17631757 |
| CG163 | Punta Corbaghiola (CG) | H1023 | H2002 | ERR17631645 |
| IZ03 | Inzecca (IZ) | H1008 | H2002 | ERR17631759 |
| IZ17 | Inzecca (IZ) | H1008 | H2002 | |
| IZ18 | Inzecca (IZ) | H1008 | H1038 | ERR17631573 |
| IZ20 | Inzecca (IZ) | H1023 | H1038 | ERR17631788 |
| IZ21 | Inzecca (IZ) | H1008 | H1097 | |
| IZ48 | Inzecca (IZ) | H1008 | H2002 | |
| IZ74 | Inzecca (IZ) | H1038 | H2002 | |
| IZ77 | Inzecca (IZ) | H1012 | H1097 | ERR17631682 |
| IZ78 | Inzecca (IZ) | H1004 | H2002 | ERR17631618 |
| IZ81 | Inzecca (IZ) | H1008 | H1012 | |
| IZ90 | Inzecca (IZ) | H1008 | H2002 | |
| IZ91 | Inzecca (IZ) | H2002 | H2002 | ERR17631627 |
| IZ92 | Inzecca (IZ) | H1038 | H2002 | |
| IZ96 | Inzecca (IZ) | H1023 | H2002 | ERR17631627 |
| IZ100 | Inzecca (IZ) | H1038 | H1097 | ERR17631736 |

| | | | | |
|---|---|---|---|---|
| IZ101 | Inzecca (IZ) | H1097 | H1063 | |
| IZ104 | Inzecca (IZ) | H1094 | H2002 | ERR17631690 |
| IZ106 | Inzecca (IZ) | H1094 | H1097 | ERR17631577 |
| IZ107 | Inzecca (IZ) | H1008 | H1097 | ERR17631624 |
| IZ108 | Inzecca (IZ) | H1063 | H1094 | ERR17631693 |
| IZ109 | Inzecca (IZ) | H1023 | H1063 | ERR17631545 |
| IZ112 | Inzecca (IZ) | H1007 | H1097 | ERR17631561 |
| IZ113 | Inzecca (IZ) | H1006 | H1097 | ERR17631758 |
| IZ114 | Inzecca (IZ) | H1007 | H1097 | ERR17631558 |
| IZ115 | Inzecca (IZ) | H1097 | H2002 | ERR17631548 |
| IZ117 | Inzecca (IZ) | H1008 | H1023 | ERR17631575 |
| IZ119 | Inzecca (IZ) | H1012 | H1094 | ERR17631720 |
| IZ121 | Inzecca (IZ) | H1012 | H1023 | ERR17631775 |
| IZ122 | Inzecca (IZ) | H1008 | H1097 | ERR17631667 |
| IZ123 | Inzecca (IZ) | H1008 | H1023 | ERR17631661 |
| IZ124 | Inzecca (IZ) | H1063 | H2002 | |
| IZ125 | Inzecca (IZ) | H1094 | H2002 | ERR17631595 |
| IZ126 | Inzecca (IZ) | H1094 | H2002 | |
| IZ127 | Inzecca (IZ) | H1008 | H1097 | ERR17631688 |
| IZ130 | Inzecca (IZ) | H1012 | H1063 | ERR17631644 |
| IZ131 | Inzecca (IZ) | H1012 | H1097 | ERR17631698 |
| IZ132 | Inzecca (IZ) | H1023 | H2002 | |
| IZ133 | Inzecca (IZ) | H1094 | H1063 | ERR17631601 |
| IZ134 | Inzecca (IZ) | H1012 | H1063 | ERR17631560 |
| IZ135 | Inzecca (IZ) | H1094 | H1063 | ERR17631703 |
| IZ136 | Inzecca (IZ) | H1008 | H1012 | ERR17631547 |
| IZ137 | Inzecca (IZ) | H1038 | H2002 | |
| IZ139 | Inzecca (IZ) | H2002 | H2002 | ERR17631568 |
| IZ140 | Inzecca (IZ) | H1008 | H2002 | |
| IZ144 | Inzecca (IZ) | H1008 | H2002 | |
| IZ147 | Inzecca (IZ) | H1008 | H1097 | ERR17631705 |
| IZ148 | Inzecca (IZ) | H1038 | H2002 | ERR17631638 |
| IZ149 | Inzecca (IZ) | H1038 | H2002 | |
| IZ150 | Inzecca (IZ) | H1023 | H2002 | |
| IZ156 | Inzecca (IZ) | H1006 | H1094 | ERR17631605 |
| IZ157 | Inzecca (IZ) | H1012 | H2002 | |
| IZ159 | Inzecca (IZ) | H2002 | H2002 | ERR17631616 |
| IZ160 | Inzecca (IZ) | H1094 | H2002 | ERR17631612 |
| IZ161 | Inzecca (IZ) | H1023 | H2002 | |

| | | | | |
|---|---|---|---|---|
| IZ162 | Inzecca (IZ) | H1008 | H1023 | ERR17631610 |
| IZ163 | Inzecca (IZ) | H1008 | H1012 | |
| IZ164 | Inzecca (IZ) | H1097 | H1063 | ERR17631639 |
| IZ167 | Inzecca (IZ) | H1008 | H1012 | |
| IZ168 | Inzecca (IZ) | H1008 | H1023 | ERR17631628 |
| IZ170 | Inzecca (IZ) | H1008 | H2002 | |
| IZ171 | Inzecca (IZ) | H2002 | H2002 | ERR17631695 |
| IZ174 | Inzecca (IZ) | H1008 | H1097 | ERR17631554 |
| IZ177 | Inzecca (IZ) | H1006 | H1094 | ERR17631735 |
| IZ178 | Inzecca (IZ) | H1006 | H1023 | ERR17631619 |
| IZ180 | Inzecca (IZ) | H1097 | H1063 | ERR17631572 |
| IZ181 | Inzecca (IZ) | H1023 | H1063 | ERR17631656 |
| IZ182 | Inzecca (IZ) | H1097 | H2002 | ERR17631747 |
| IZ183 | Inzecca (IZ) | H1023 | H1097 | ERR17631562 |
| IZ184 | Inzecca (IZ) | H1006 | H1008 | ERR17631726 |
| IZ186 | Inzecca (IZ) | H1006 | H1023 | ERR17631660 |
| IZ187 | Inzecca (IZ) | H1006 | H1023 | ERR17631604 |
| IZ188 | Inzecca (IZ) | H1023 | H2002 | ERR17631659 |
| IZ189 | Inzecca (IZ) | H1008 | H2002 | ERR17631651 |
| IZ190 | Inzecca (IZ) | H1094 | H1023 | ERR17631700 |
| TG_A56 | Teghime zone A (TG_A) | H1045 | H2002 | |
| TG_A57 | Teghime zone A (TG_A) | H1045 | H1092 | ERR17631724 |
| TG_A60 | Teghime zone A (TG_A) | H1016 | H2002 | |
| TG_A61 | Teghime zone A (TG_A) | H1016 | H2002 | |
| TG_A62 | Teghime zone A (TG_A) | H1016 | H2002 | |
| TG_A63 | Teghime zone A (TG_A) | H1028 | H1045 | ERR17631673 |
| TG_A64 | Teghime zone A (TG_A) | H1016 | H1028 | ERR17631676 |
| TG_A65 | Teghime zone A (TG_A) | H1028 | H1092 | ERR17631753 |
| TG_A66 | Teghime zone A (TG_A) | H1028 | H1092 | ERR17631576 |
| TG_A67 | Teghime zone A (TG_A) | H1028 | H1092 | ERR17631543 |
| TG_A68 | Teghime zone A (TG_A) | H1028 | H1045 | ERR17631740 |
| TG_A69 | Teghime zone A (TG_A) | H1028 | H1045 | ERR17631699 |
| TG_A70 | Teghime zone A (TG_A) | H1028 | H1045 | ERR17631588 |
| TG_A71 | Teghime zone A (TG_A) | H1028 | H1045 | ERR17631549 |
| TG_A74 | Teghime zone A (TG_A) | H1028 | H1045 | ERR17631672 |
| TG_A75 | Teghime zone A (TG_A) | H1028 | H1045 | ERR17631783 |
| TG_A76 | Teghime zone A (TG_A) | H1028 | H1045 | ERR17631579 |
| TG_A78 | Teghime zone A (TG_A) | H1028 | H1045 | ERR17631692 |
| TG_A79 | Teghime zone A (TG_A) | H1028 | H1045 | ERR17631550 |

| | | | | |
|---|---|---|---|---|
| TG_A80 | Teghime zone A (TG_A) | H1028 | H1045 | ERR17631710 |
| TG_A82 | Teghime zone A (TG_A) | H1028 | H1045 | ERR17631606 |
| TG_A83 | Teghime zone A (TG_A) | H1028 | H1045 | ERR17631632 |
| TG_A84 | Teghime zone A (TG_A) | H1028 | H1045 | ERR17631608 |
| TG_A85 | Teghime zone A (TG_A) | H1028 | H1045 | ERR17631750 |
| TG_A86 | Teghime zone A (TG_A) | H1045 | H1092 | ERR17631718 |
| TG_A87 | Teghime zone A (TG_A) | H1028 | H1063 | ERR17631655 |
| TG_A88 | Teghime zone A (TG_A) | H1045 | H1092 | ERR17631666 |
| TG_A89 | Teghime zone A (TG_A) | H1028 | H1045 | ERR17631709 |
| TG_A91 | Teghime zone A (TG_A) | H1016 | H1092 | ERR17631641 |
| TG_A93 | Teghime zone A (TG_A) | H1028 | H1063 | ERR17631708 |
| TG_A94 | Teghime zone A (TG_A) | H1028 | H1063 | ERR17631654 |
| TG_A95 | Teghime zone A (TG_A) | H1028 | H1063 | ERR17631744 |
| TG_A96 | Teghime zone A (TG_A) | H1028 | H1063 | ERR17631768 |
| TG_A103 | Teghime zone A (TG_A) | H1016 | H1028 | ERR17631772 |
| TG_A104 | Teghime zone A (TG_A) | H1092 | H1093 | ERR17631646 |
| TG_A106 | Teghime zone A (TG_A) | H1028 | H1063 | ERR17631617 |
| TG_A305 | Teghime zone A (TG_A) | H1028 | H2002 | ERR17631636 |
| TG_A306 | Teghime zone A (TG_A) | H1028 | H2002 | ERR17631704 |
| TG_A307 | Teghime zone A (TG_A) | H1028 | H1045 | ERR17631581 |
| TG_A308 | Teghime zone A (TG_A) | H1045 | H1092 | ERR17631745 |
| TG_A309 | Teghime zone A (TG_A) | H1045 | H1092 | ERR17631729 |
| TG_A317 | Teghime zone A (TG_A) | H1028 | H1092 | ERR17631643 |
| TG_A319 | Teghime zone A (TG_A) | H1028 | H1045 | ERR17631546 |
| TG_A320 | Teghime zone A (TG_A) | H1028 | H1045 | ERR17631773 |
| TG_A321 | Teghime zone A (TG_A) | H1028 | H1063 | ERR17631631 |
| TG_A322 | Teghime zone A (TG_A) | H1028 | H1045 | ERR17631684 |
| TG_A324 | Teghime zone A (TG_A) | H1028 | H1045 | ERR17631780 |
| TG_A326 | Teghime zone A (TG_A) | H1028 | H1063 | ERR17631571 |
| TG_A327 | Teghime zone A (TG_A) | H1045 | H1092 | ERR17631742 |
| TG_A342 | Teghime zone A (TG_A) | H1028 | H1045 | ERR17631782 |
| TG_A343 | Teghime zone A (TG_A) | H1028 | H1045 | ERR17631587 |
| TG_A346 | Teghime zone A (TG_A) | H1045 | H1092 | ERR17631785 |
| TG_A347 | Teghime zone A (TG_A) | H1045 | H1092 | ERR17631540 |
| TG_B11 | Teghime zone B (TG_B) | H1028 | H1045 | ERR17631603 |
| TG_B15 | Teghime zone B (TG_B) | H1052 | H2002 | ERR17631707 |
| TG_B33 | Teghime zone B (TG_B) | H1013 | H2002 | |
| TG_B35 | Teghime zone B (TG_B) | H1013 | H2002 | |
| TG_B107 | Teghime zone B (TG_B) | H1016 | H1097 | ERR17631634 |

| | | | | |
|---|---|---|---|---|
| TG_B108 | Teghime zone B (TG_B) | H1102 | H2015 | ERR17631732 |
| TG_B110 | Teghime zone B (TG_B) | H1016 | H1068 | |
| TG_B111 | Teghime zone B (TG_B) | H1016 | H1033 | ERR17631629 |
| TG_B112 | Teghime zone B (TG_B) | H1036 | H1052 | ERR17631623 |
| TG_B113 | Teghime zone B (TG_B) | H1036 | H1052 | ERR17631541 |
| TG_B114 | Teghime zone B (TG_B) | H1013 | H1036 | ERR17631697 |
| TG_B115 | Teghime zone B (TG_B) | H1016 | H1033 | ERR17631680 |
| TG_B121 | Teghime zone B (TG_B) | H1036 | H1052 | ERR17631585 |
| TG_B122 | Teghime zone B (TG_B) | H1045 | H2015 | ERR17631770 |
| TG_B123 | Teghime zone B (TG_B) | H1013 | H1052 | ERR17631752 |
| TG_B124 | Teghime zone B (TG_B) | H1052 | H1068 | ERR17631717 |
| TG_B126 | Teghime zone B (TG_B) | H1028 | H1052 | ERR17631683 |
| TG_B127 | Teghime zone B (TG_B) | H1028 | H1036 | ERR17631566 |
| TG_B132 | Teghime zone B (TG_B) | H1036 | H1097 | ERR17631633 |
| TG_B135 | Teghime zone B (TG_B) | H1016 | H1085 | ERR17631599 |
| TG_B136 | Teghime zone B (TG_B) | H1068 | H2002 | |
| TG_B138 | Teghime zone B (TG_B) | H1016 | H1068 | |
| TG_B158 | Teghime zone B (TG_B) | H1016 | H1036 | |
| TG_B159 | Teghime zone B (TG_B) | H1016 | H1036 | |
| TG_B160 | Teghime zone B (TG_B) | H1036 | H2002 | |
| TG_B161 | Teghime zone B (TG_B) | H1016 | H1036 | ERR17631765 |
| TG_B163 | Teghime zone B (TG_B) | H1016 | H1036 | ERR17631774 |
| TG_B165 | Teghime zone B (TG_B) | H1036 | H1068 | ERR17631578 |
| TG_B168 | Teghime zone B (TG_B) | H1016 | H1068 | |
| TG_B170 | Teghime zone B (TG_B) | H1016 | H2005 | ERR17631536 |
| TG_B172 | Teghime zone B (TG_B) | H1016 | H1068 | |
| TG_B173 | Teghime zone B (TG_B) | H1016 | H2005 | |
| TG_B174 | Teghime zone B (TG_B) | H1016 | H2005 | ERR17631555 |
| TG_B179 | Teghime zone B (TG_B) | H1013 | H1052 | ERR17631622 |
| TG_B185 | Teghime zone B (TG_B) | H1068 | H2005 | ERR17631691 |
| TG_B186 | Teghime zone B (TG_B) | H1013 | H1016 | ERR17631662 |
| TG_B189 | Teghime zone B (TG_B) | H1016 | H2015 | ERR17631589 |
| TG_B190 | Teghime zone B (TG_B) | H1028 | H1068 | ERR17631611 |
| TG_B192 | Teghime zone B (TG_B) | H1013 | H1016 | ERR17631590 |
| TG_B196 | Teghime zone B (TG_B) | H1068 | H2002 | |
| TG_B204 | Teghime zone B (TG_B) | H1013 | H2005 | ERR17631553 |
| TG_B205 | Teghime zone B (TG_B) | H1013 | H1052 | ERR17631615 |
| TG_B206 | Teghime zone B (TG_B) | H1013 | H1068 | ERR17631741 |
| TG_B207 | Teghime zone B (TG_B) | H1036 | H1045 | ERR17631763 |

| | | | | |
|---|---|---|---|---|
| TG_B208 | Teghime zone B (TG_B) | H1013 | H1036 | ERR17631778 |
| TG_B209 | Teghime zone B (TG_B) | H1028 | H1036 | ERR17631743 |
| TG_B212 | Teghime zone B (TG_B) | H1013 | H2005 | ERR17631614 |
| TG_B215 | Teghime zone B (TG_B) | H1068 | H2005 | ERR17631563 |
| TG_B216 | Teghime zone B (TG_B) | H1016 | H1068 | ERR17631534 |
| TG_B217 | Teghime zone B (TG_B) | H1016 | H1052 | ERR17631607 |
| TG_B219 | Teghime zone B (TG_B) | H1068 | H2005 | ERR17631653 |
| TG_B220 | Teghime zone B (TG_B) | H1013 | H1068 | ERR17631557 |
| TG_B221 | Teghime zone B (TG_B) | H1068 | H2005 | ERR17631584 |
| TG_B223 | Teghime zone B (TG_B) | H1045 | H2005 | ERR17631685 |
| TG_B224 | Teghime zone B (TG_B) | H1013 | H1068 | ERR17631658 |
| TG_B228 | Teghime zone B (TG_B) | H1013 | H2002 | |
| TG_B230 | Teghime zone B (TG_B) | H1013 | H2005 | ERR17631674 |
| TG_B233 | Teghime zone B (TG_B) | H1013 | H1016 | ERR17631583 |
| TG_B234 | Teghime zone B (TG_B) | H1013 | H2002 | |
| TG_B238 | Teghime zone B (TG_B) | H1045 | H1097 | ERR17631602 |
| TG_B240 | Teghime zone B (TG_B) | H1013 | H1068 | ERR17631597 |
| TG_B241 | Teghime zone B (TG_B) | H1016 | H2005 | ERR17631749 |
| TG_B242 | Teghime zone B (TG_B) | H1016 | H2005 | ERR17631781 |
| TG_B243 | Teghime zone B (TG_B) | H1016 | H2005 | ERR17631620 |
| TG_B244 | Teghime zone B (TG_B) | H1016 | H2005 | ERR17631723 |
| TG_B248 | Teghime zone B (TG_B) | H1052 | H2002 | ERR17631762 |
| TG_B253 | Teghime zone B (TG_B) | H1068 | H2005 | ERR17631542 |
| TG_B258 | Teghime zone B (TG_B) | H1013 | H1068 | ERR17631570 |
| TG_B259 | Teghime zone B (TG_B) | H1013 | H1068 | ERR17631635 |
| TG_B260 | Teghime zone B (TG_B) | H1013 | H1068 | ERR17631733 |
| TG_B262 | Teghime zone B (TG_B) | H2002 | H2005 | ERR17631771 |
| TG_B263 | Teghime zone B (TG_B) | H2002 | H2005 | ERR17631760 |
| TG_B264 | Teghime zone B (TG_B) | H2002 | H2005 | ERR17631565 |
| TG_B267 | Teghime zone B (TG_B) | H1016 | H2005 | ERR17631671 |
| TG_B271 | Teghime zone B (TG_B) | H1016 | H2005 | |
| TG_B272 | Teghime zone B (TG_B) | H1016 | H2005 | ERR17631787 |
| TG_B273 | Teghime zone B (TG_B) | H1016 | H2005 | ERR17631722 |
| TG_B274 | Teghime zone B (TG_B) | H1016 | H2005 | ERR17631766 |
| TG_B275 | Teghime zone B (TG_B) | H1016 | H2005 | ERR17631613 |
| TG_B276 | Teghime zone B (TG_B) | H1028 | H2002 | ERR17631739 |
| TG_B278 | Teghime zone B (TG_B) | H2002 | H2005 | ERR17631591 |
| TG_B283 | Teghime zone B (TG_B) | H1068 | H1036 | ERR17631642 |
| TG_B285 | Teghime zone B (TG_B) | H1036 | H2005 | ERR17631609 |

| | | | | |
|---|---|---|---|---|
| TG_B286 | Teghime zone B (TG_B) | H1028 | H2002 | ERR17631721 |
| TG_B289 | Teghime zone B (TG_B) | H1028 | H2002 | ERR17631552 |
| TG_B291 | Teghime zone B (TG_B) | H1028 | H2002 | ERR17631679 |
| TG_B292 | Teghime zone B (TG_B) | H1013 | H2005 | ERR17631596 |
| TG_B293 | Teghime zone B (TG_B) | H1016 | H2005 | ERR17631663 |
| TG_B296 | Teghime zone B (TG_B) | H1028 | H1036 | ERR17631593 |
| TG_B297 | Teghime zone B (TG_B) | H1028 | H1036 | ERR17631600 |
| TG_B298 | Teghime zone B (TG_B) | H1013 | H1045 | |
| TG_B299 | Teghime zone B (TG_B) | H1068 | H2002 | ERR17631580 |
| TG_B300 | Teghime zone B (TG_B) | H1016 | H1068 | |
| TG_B301 | Teghime zone B (TG_B) | H1004 | H2005 | ERR17631746 |
| TG_B302 | Teghime zone B (TG_B) | H1028 | H2005 | ERR17631748 |
| TG_B303 | Teghime zone B (TG_B) | H1068 | H2005 | ERR17631730 |
| TG_B304 | Teghime zone B (TG_B) | H2002 | H2005 | ERR17631754 |
| TG_B329 | Teghime zone B (TG_B) | H1045 | H2015 | ERR17631764 |
| TG_B332 | Teghime zone B (TG_B) | H2002 | H2005 | ERR17631694 |
| TG_B335 | Teghime zone B (TG_B) | H1004 | H2005 | ERR17631640 |
| TG_B340 | Teghime zone B (TG_B) | H1016 | H2005 | ERR17631731 |
